# Meta-transcriptomic detection of diverse and divergent RNA viruses in green and chlorarachniophyte algae

**DOI:** 10.1101/2020.06.08.141184

**Authors:** Justine Charon, Vanessa Rossetto Marcelino, Richard Wetherbee, Heroen Verbruggen, Edward C. Holmes

## Abstract

Our knowledge of the diversity and evolution of the virosphere will likely increase dramatically with the study of microbial eukaryotes, including the microalgae in few RNA viruses have been documented to date. By combining meta-transcriptomic approaches with sequence and structural-based homology detection, followed by PCR confirmation, we identified 18 novel RNA viruses in two major groups of microbial algae – the chlorophytes and the chlorarachniophytes. Most of the RNA viruses identified in the green algae class Ulvophyceae were related to those from the families *Tombusviridae* and *Amalgaviridae* that have previously been associated with plants, suggesting that these viruses have an evolutionary history that extends to when their host groups shared a common ancestor. In contrast, seven ulvophyte associated viruses exhibited clear similarity with the mitoviruses that are most commonly found in fungi. This is compatible with horizontal virus transfer between algae and fungi, although mitoviruses have recently been documented in plants. We also document, for the first time, RNA viruses in the chlorarachniophytes, including the first observation of a negative-sense (bunya-like) RNA virus in microalgae. The other virus-like sequence detected in chlorarachniophytes is distantly related to those from the plant virus family *Virgaviridae*, suggesting that they may have been inherited from the secondary chloroplast endosymbiosis event that marked the origin of the chlorarachniophytes. More broadly, this work suggests that the scarcity of RNA viruses in algae most likely results from limited investigation rather than their absence. Greater effort is needed to characterize the RNA viromes of unicellular eukaryotes, including through structure-based methods that are able to detect distant homologies, and with the inclusion of a wider range of eukaryotic microorganisms.

**Author summary:** RNA viruses are expected to infect all living organisms on Earth. Despite recent developments in and the deployment of large-scale sequencing technologies, our understanding of the RNA virosphere remains anthropocentric and largely restricted to human, livestock, cultivated plants and vectors for viral disease. However, a broader investigation of the diversity of RNA viruses, especially in protists, is expected to answer fundamental questions about their origin and long-term evolution. This study first investigates the RNA virus diversity in unicellular algae taxa from the phylogenetically distinct ulvophytes and chlorarachniophytes taxa. Despite very high levels of sequence divergence, we were able to identify 18 new RNA viruses, largely related to plant and fungi viruses, and likely illustrating a past history of horizontal transfer events that have occurred during RNA virus evolution. We also hypothesise that the sequence similarity between a chlorarachniophyte-associated virga-like virus and members of *Virgaviridae* associated with plants may represent inheritance from a secondary endosymbiosis event. A promising approach to detect the signals of distant virus homologies through the analysis of protein structures was also utilised, enabling us to identify potential highly divergent algal RNA viruses.

## INTRODUCTION

Viruses are likely to infect every cellular organism and play fundamental roles in biosphere diversity, evolution, and ecology. Studies of the global virosphere performed to date have revealed marked heterogeneities in virus composition. For example, while RNA viruses are commonplace in eukaryotes, they are less often found in bacteria, with only two families described to date, and are yet to be conclusively identified in Archaea. Rather, both the bacteria and Archaea are dominated by DNA viruses^1,2^. It is unclear, however, whether the highly skewed distribution of viruses reflects fundamental biological, cellular or ecological factors of the hosts in question, or because RNA viruses in bacteria and Archaea are often so divergent in sequence that they are difficult to detect using primary sequence comparisons alone.

Despite greatly increased virus sampling following the advent of metagenomic next-generation sequencing, our picture of the virosphere remains largely restricted to bacteria, vertebrates and plants^3,4^. Clearly, such a sampling bias will also impact our knowledge of the fundamental patterns and processes of virus origins and evolution. A good example of this major knowledge bias are the unicellular eukaryotes, grouped under the term “protists”, and particularly the microalgae: only 61 distinct viruses have been formally recognized in these taxa^5^, with only 82 viral sequences classified as infecting eukaryotic microalgae^6^. This represents only 0.6% of the total 14,679 viral sequences listed on the Viral-Host database (release April 2020), although the true number of microalgal species is estimated to exceed 300,000^7^.

Despite early attempts, and the first algal virus cultivation in 1979^8,9^, the isolation and characterization of phycoviruses (i.e. algal viruses) has been constrained by the difficulty in cultivating both the algae and their viruses, as well as the inherent challenges in identifying RNA viruses that are highly divergent in primary sequence. Indeed, because RNA viruses are the fastest evolving entities described, phylogenetic signal is rapidly lost over evolutionary time. Hence, it is possible that the low number of algal RNA viruses detected to date simply reflects the fact that they are highly divergent in sequence, even in the canonical RNA-dependent RNA polymerase (RdRp), and hence refractory to detection using primary sequence similarity. Importantly, protein structures are expected to be an order of magnitude more conserved than amino acid sequences^10^. As a consequence, the study of conserved secondary or tertiary structures could help identify distant homologies among RNA viruses over extended evolutionary time-scales^11,12^. Accordingly, the analysis of conserved protein structures may constitute a promising approach to identify novel viruses within the microalgae.

Microalgae are a polyphyletic group of microscopic unicellular photosynthetic organisms distributed across diverse branches of the eukaryotic phylogeny. With 72,500 species discovered to date and organised into the TSAR (Telonemia, Stramenopiles, Alveolates, Rhizaria), Archaeplastida, Haptista, Cryptista and “excavates” supergroups^13–15^, microalgae constitute a huge source of genetic diversity^16^. In addition to their wide range of habitats, their ubiquity and the diversity of genomic features characterised to date (with linear, circular, segmented, non-segmented, single-strand and double-strand genomes) suggest that microalgae will harbour an enormous untapped source of viral diversity. Due to the ancient nature of eukaryotic algae (ca. 1.8 billion years), their involvement in secondary plastid endosymbiosis events involving many branches of the eukaryotic phylogeny, and that microalgae constitute a primary food source for marine and freshwater food chain, it is also possible that algal viruses played a crucial role in the early events of eukaryote virus evolution^5,17^.

The algal viruses documented to date are dominated by those with DNA genomes: 55 of the 82 algal virus sequences available at VirusHostdb. These include the well-known giant viruses, the majority of which (53%) have been described in the green algae (Chlorophyta). This DNA-dominated virome of green algae contrasts with those of their sister-group, the land plants^18^, for which 60% of the 3590 viral sequences are RNA viruses (i.e. the “Riboviria”; VirusHostdb, April 2020 release). In marked contrast, 85% of the 27 algae RNA viruses described to date have been identified in diatoms^19^. Even the few algal RNA viruses characterised to date display impressive diversity, belonging to the families *Totiviridae*, *Reoviridae*, *Marnaviridae*, *Endornaviridae*, *Flaviviridae*, *Narnaviridae* and *Alvernaviridae*.

However, it is currently unclear whether these seemingly differing distributions of DNA and RNA viruses reflect a major switch in virus composition that occurred during the expansion of land plants or is simply indicative of the inherent limitations in sampling and investigation described above^20^. As such, revealing the pattern and extent of RNA virus diversity in algae may have profound consequences for understanding the long-term processes that have shaped virus diversity and evolution.

Herein, we characterised more of the RNA virosphere in cultured samples of two clades of microalgae: (i) the green algae (Chlorophyta), that are part of the Archaeplastida eukaryotic supergroup, and (ii) the chlorarachniophytes, a lineage of Rhizaria that obtained a chloroplast through secondary endosymbiosis of a green alga^21^. To this end we performed an unbiased (i.e. bulk RNA-sequencing) meta-transcriptomic analysis with an emphasis on detecting remote signals of homology in the RdRp, the hallmark of RNA viruses, using protein-profile based approaches. The comparison of the viromes of these two distant groups will help answer the following key questions: (i) is the long-term divergence between the two algae taxa also reflected in their RNA virome compositions? (ii) does their RNA virome provide evidence for complex evolutionary histories, including horizontal transfer events? (iii) is the RNA/DNA virus bias observed between algae and green plants an artefact of sampling or reflect a more fundamental biological division? More generally, we aimed to broaden our understanding of the biodiversity of algal viruses, with the expectation that the characterization of the algal virosphere will have important implications for understanding and managing the roles they play in global element cycling, climate forcing and biotechnology. Indeed, viruses are important in regulating microalgae populations and may provide a key reservoir for genetic novelty^22,23^.

## RESULTS

### The RNA viromes of two divergent groups of microalgae

Our aim was to determine the RNA viromes of six microalgae cultures from six different algal species classified into two highly phylogenetically distinct algal clades: the chlorarachniophytes (Rhizaria) and the green algae (Chlorophyta, Archaeplastida) (Fig 1A). To the best of our knowledge, this is the first identification of viruses in the Chlorarachniophyceae and Ulvophyceae (Chlorophyta) classes (Fig 1C) ^24^.

**Fig 1.**
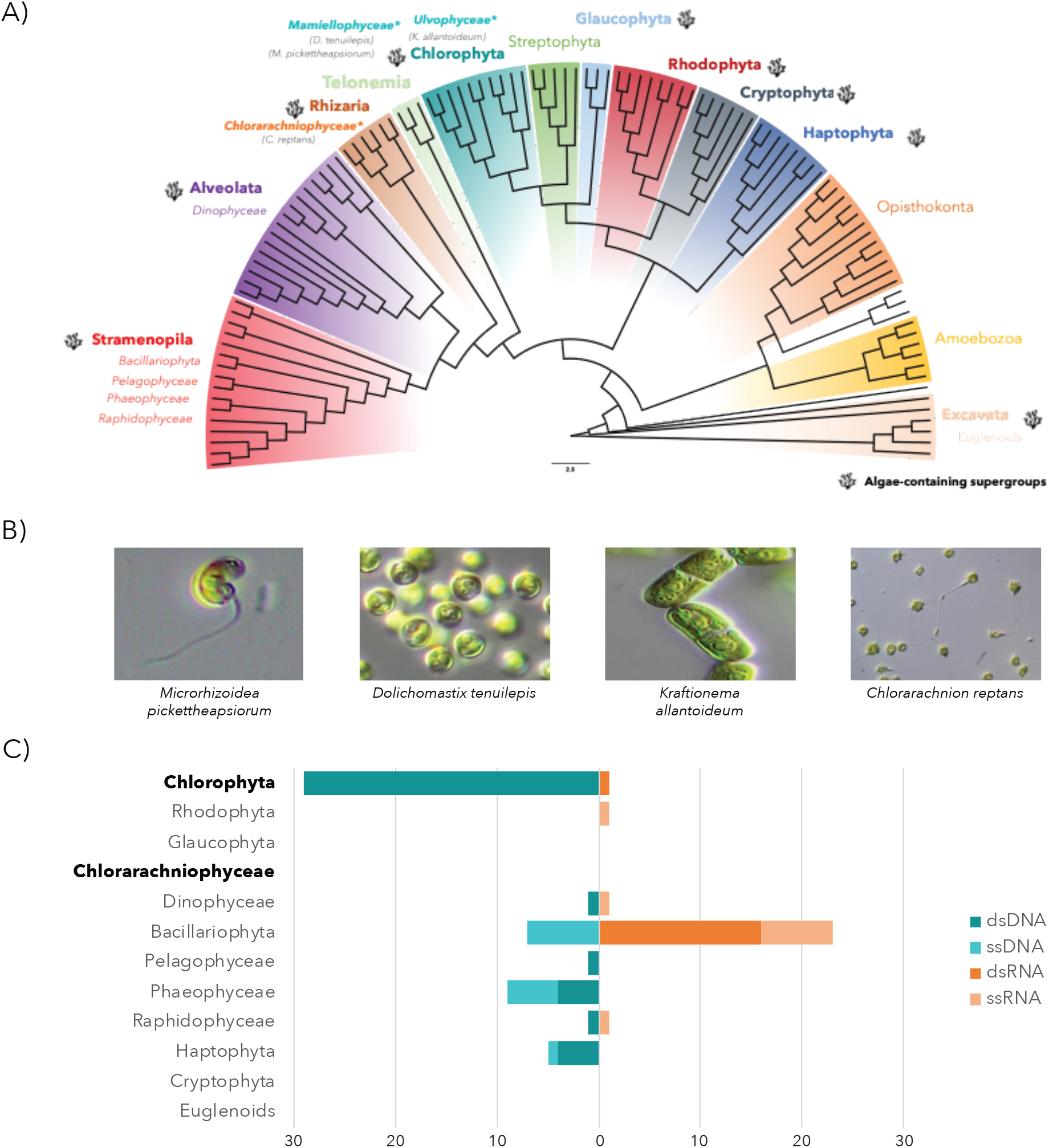
The enormous diversity of algae contrasts with its poorly-characterized virosphere. (A) Representation of algae supergroups among the diversity of eukaryotes (latest eukaryotic classification retrieved from^15^). The phylogenetic tree was adapted from^77^. Pictures illustrate some of the samples used in this study and corresponding clades are marked with “*”. (B) Pictures of algae cultures used in this study. (C) The current extent of the microalgae virosphere. The viral sequence counts for each virus class (DNA or RNA, single-stranded or double-stranded) were retrieved from VirusHostdb^6^ according to 11 major eukaryotic algae lineages. Microalgal lineages investigated in this study are highlighted in bold.

While our previous limited understanding of viruses infecting chlorarachniophytes can be explained by the small number of species from this clade characterized to date (only 15 in Algaebase), the Ulvophyceae is an abundant and diverse algal lineage in existence since the late Proterozoic and comprises at least 1,933 species^25^. It contains a wide range of morphologies from unicellular benthic algae to large seaweeds^26^ and its representatives commonly occur in marine, terrestrial and freshwater habitats^18^. We therefore performed RNA sequencing (meta-transcriptomics) on six microalgal species belonging to both the green algae (classes Ulvophyceae and Mamiellophyceae) and chlorarachniophytes (Table 1).

**Table 1.** Sample and library description.

| Library | Species | Class |
| --- | --- | --- |
| ALG_1 | *Kraftionema allantoideum* | Ulvophyceae/Kraftionemaceae |
|  | *Microrhizoidea pickettheapsiorum* | Mamiellophyceae/Dolichomastigaceae |
|  | *Dolichomastix tenuilepis* | Mamiellophyceae/Dolichomastigaceae |
| ALG_2 | *Ostreobium* sp. HV05007bc | Ulvophyceae/Bryopsidales |

| ALG_3 | *Chlorarachnion reptans* | Chlorarachniophyceae/Chlorarachnion |
| --- | --- | --- |
|  | *Lotharella* sp | Chlorarachniophyceae/Lotharella |

Because of variable levels of RNA quality and quantity, fewer non-rRNA reads were obtained for the ALG_1 and ALG_3 libraries, which also likely explains the reduced average length of contigs in these libraries compared to ALG_2.

In total, we identified 18 new putative viral sequences using a standard sequence similarity search among the three libraries, largely comprising those with double-stranded (ds) RNA or single-stranded positive-sense (ssRNA(+)) genomes (Table 2). A divergent bunya-like partial sequence was also retrieved from *Chlorarachnion reptans* and may constitute the first negative-sense (ssRNA(-)) virus identified in microalgae. Importantly, the presence of these viruses was validated by RT-PCR on each total extracted RNA (Fig S1). In each case these viruses exhibited very low levels of sequence similarity to existing RdRp amino acid sequences, with sequence identities ranging from only 27 to 38%. Most of the viral sequences discovered in this study likely correspond to full-length genomes, given that the length and genomic organization (ORF numbers, length, etc) is similar to the well-annotated reference homologs. Thus, these sequences are very likely true viruses rather than endogenous viral elements (EVEs) inserted into host genomes.

**Table 2.**
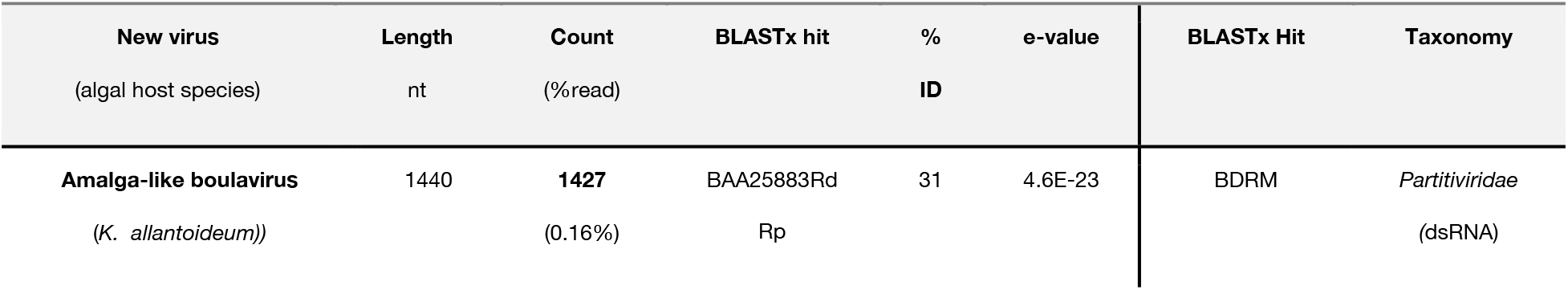
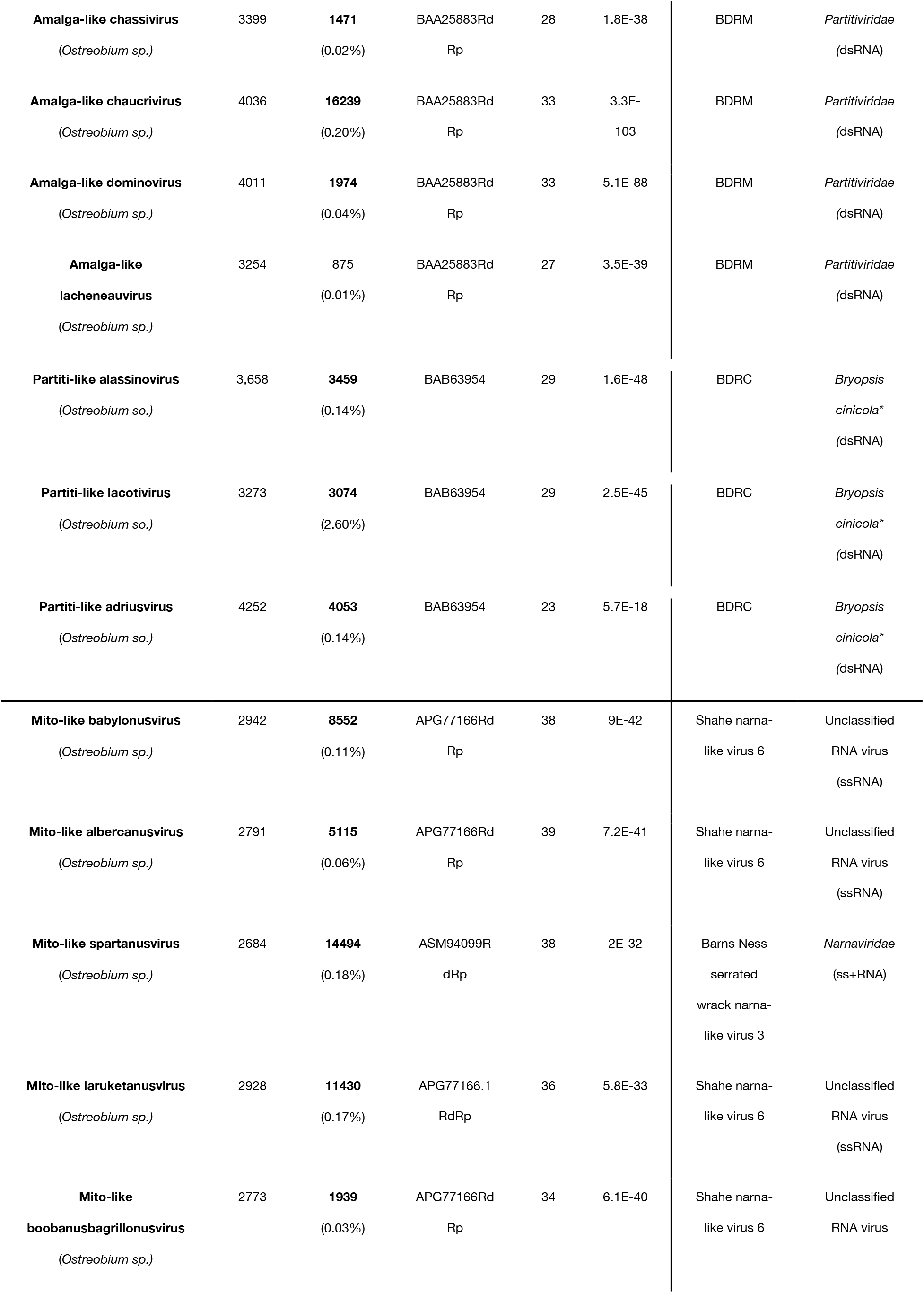
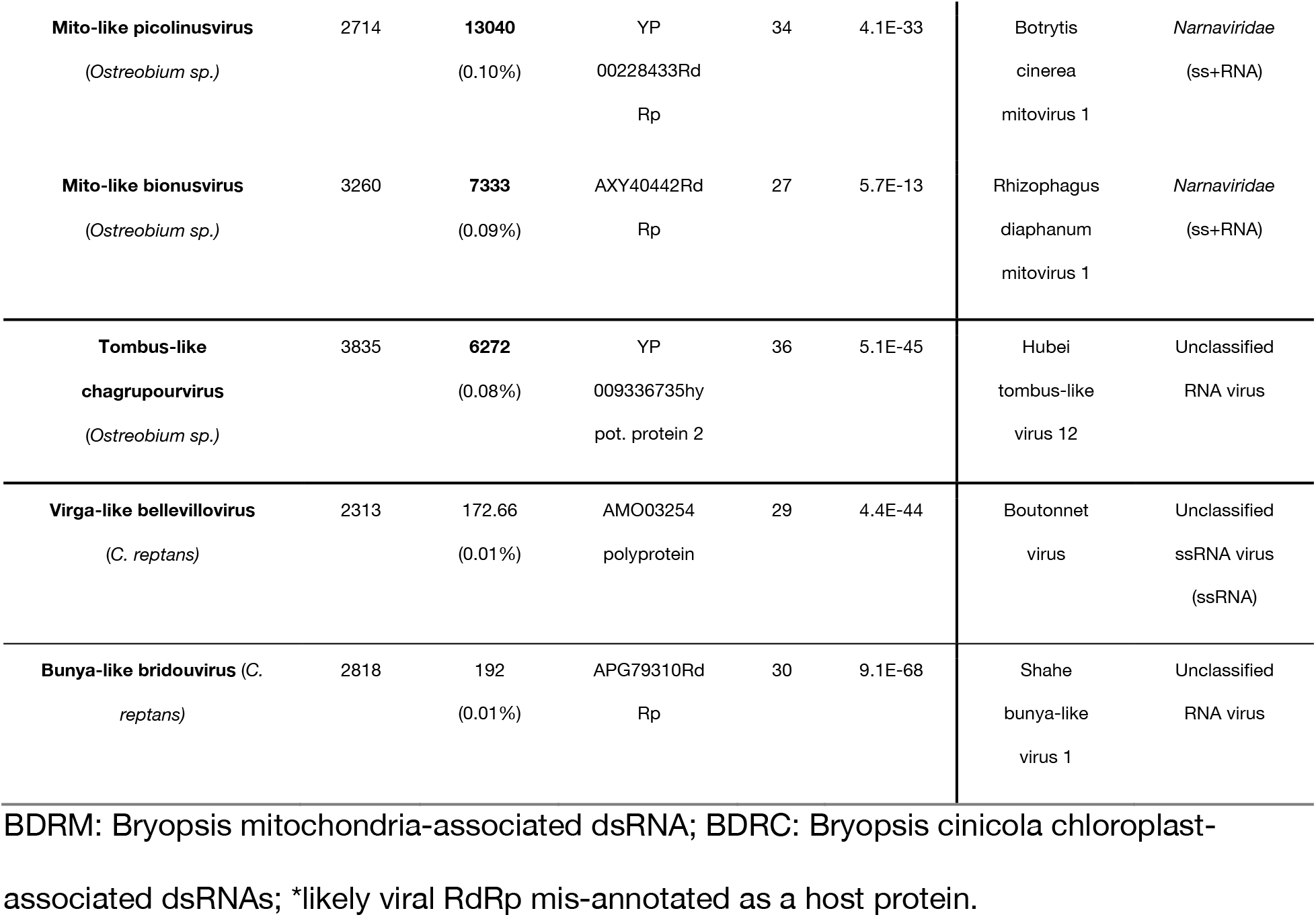
BLASTx results of the newly described virus-like sequences against nr database.

Eight of the viruses identified in *Ostreobium* sp. fell within the *Narnaviridae* or *Tombusviridae.* In contrast, five viral sequences from *Ostreobium* sp. and *K. allantoideum* do not fit into defined taxonomic groups, and were instead related to the broad set of ‘partiti-like’ viruses that comprise the *Partitiviridae, Totiviridae* and *Amalgaviridae*. Finally, more divergent but detectable sequence similarities to the *Virgaviridae* (+ssRNA) were obtained for samples from the chlorarachniophyte library.

### Detection of divergent viruses using protein structural data

An additional attempt to detect even more divergent RNA viruses was conducted was using protein structure. In particular, it is possible that highly divergent viruses are part of the unknown sequences (i.e. contigs with no match in nt/nr databases, or the ‘dark matter’) that comprise between 50-60% of total contigs obtained in this study (Fig 2A). Accordingly, we attempted to detect evolutionary-conserved features of protein structural and functional motifs in orphan sequences that encode unknown ORFs with a minimum of 200 amino acids (600 nucleotides) (Fig 2A). The corresponding translated ORFs were submitted to the protein-profile based program HMMer3 and compared with profiles from the PFAM RdRp clan and the VOG databases. To exclude false-positives, all positive hits were compared to the entire PFAM database. This resulted in the identification of three non-phage contigs that displayed RdRp homology: ALG_2_DN19089, ALG_2_DN594 and ALG_3_DN34624 (Table 3).

**Fig 2.**
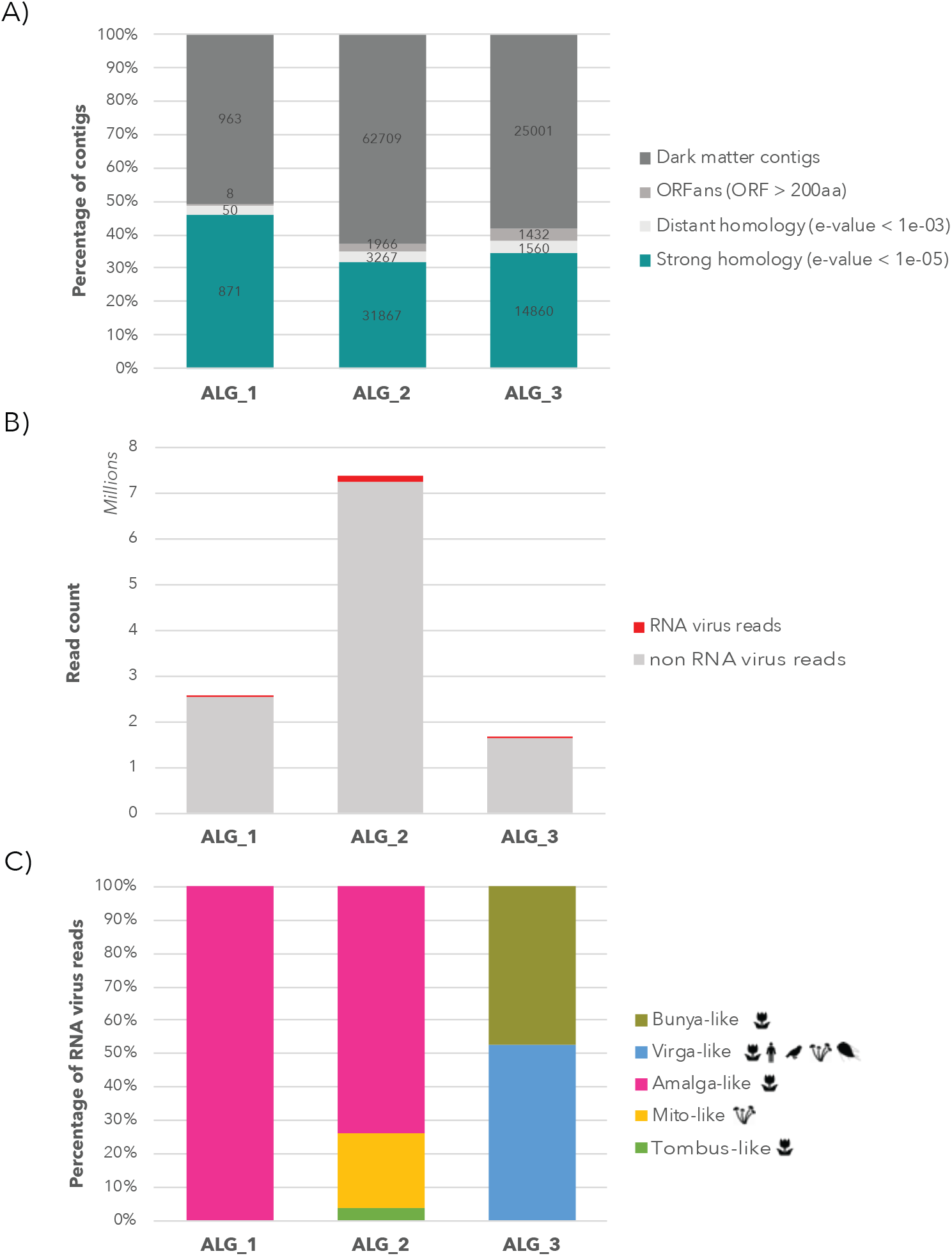
Abundance of unknown and RNA virus-like contigs detected in the algal libraries. (A) Percentage of non-assigned contigs. For clarity, numbers are normalized as the percentage of total contigs (actual contig numbers are indicated in bold). Blue: number of contigs showing a strong sequence similarity with the nr database (e-value < 1e-05); light grey: contigs showing a weak sequence similarity to the nr database (e-values 1e-05 to 1e-03); middle-dark grey: contigs with no sequence similarity detected by BLASTx/BLASTp but encoding one or more ORFs of more than 200 amino acids (600 nt); dark grey: genomic ‘dark matter’ – contigs without any signal detected or any long ORFs encoded. (B) Total number of RNA virus reads per total number of non-rRNA reads in each library. (C) Distribution of RNA virus diversity in the three libraries and percentage of RNA virus reads associated with each viral super-clade. The host range is represented for each viral clade.

**Table 3.**
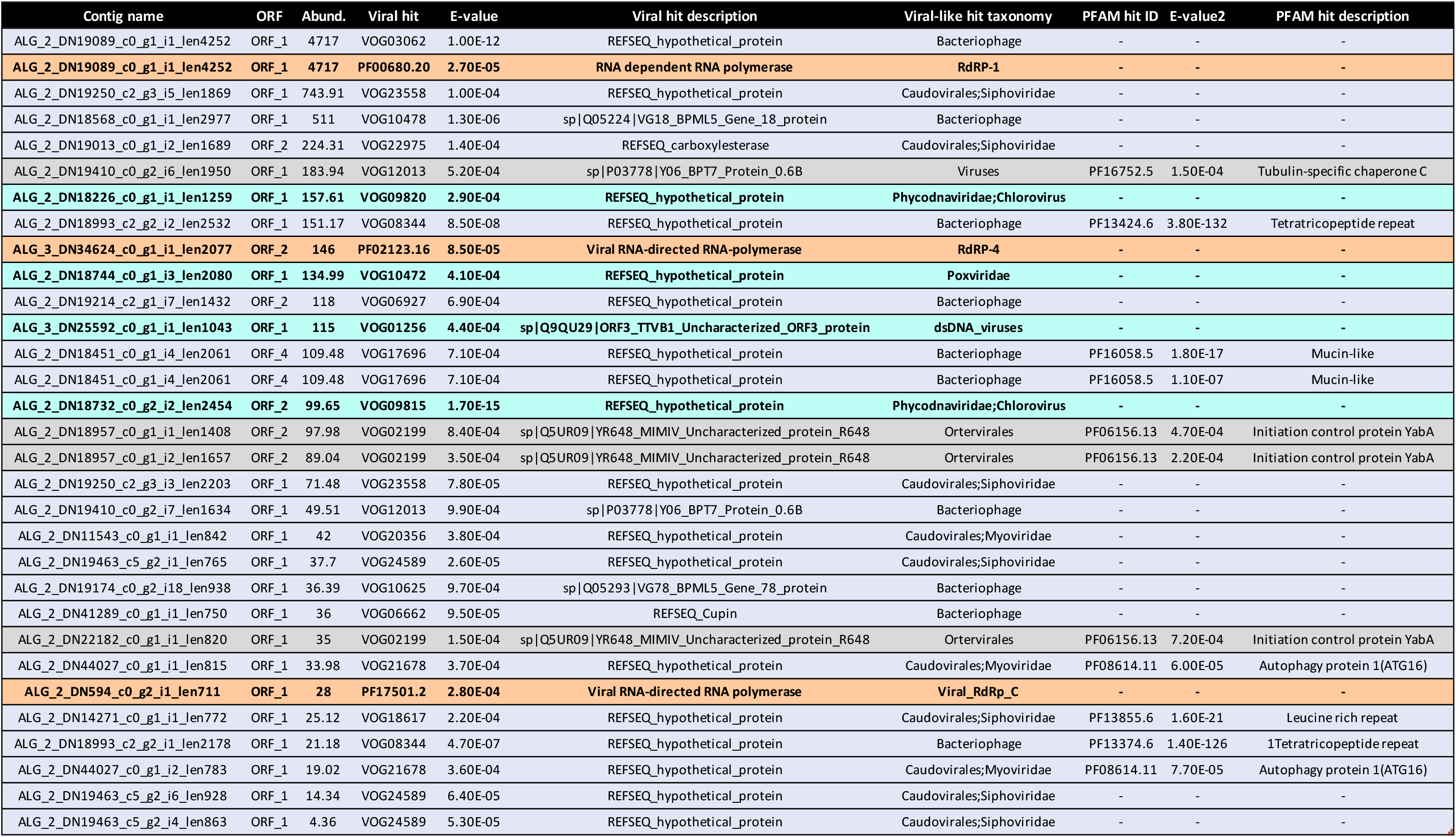
Results of the VOGdb and PFAM HMM analysis. Light purple: phage-like sequences. Grey: Non-viral sequences; Light blue: DNA virus-like sequences; Orange: RNA virus-like sequences.

| Contig name | ORF | Abund. | Viral hit | E-value | Viral hit description | Viral-like hit taxonomy | PFAM hit ID | E-value2 | PFAM hit description |
| --- | --- | --- | --- | --- | --- | --- | --- | --- | --- |
| ALG_2_DN19089_c0_g1_i1_len4252 | ORF_1 | 4717 | VOG03062 | 1.00E-12 | REFSEQ_hypothetical_protein | Bacteriophage | - | - | - |
| **ALG_2_DN19089_c0_g1_i1_len4252** | **ORF_1** | **4717** | **PF00680.20** | **2.70E-05** | **RNA dependent RNA polymerase** | **RdRP-1** | - | - | - |
| ALG_2_DN19250_c2_g3_i5_len1869 | ORF_1 | 743.91 | VOG23558 | 1.00E-04 | REFSEQ_hypothetical_protein | Caudovirales;Siphoviridae | - | - | - |
| ALG_2_DN18568_c0_g1_i1_len2977 | ORF_1 | 511 | VOG10478 | 1.30E-06 | sp\|Q05224\|VG18_BPML5_Gene_18_protein | Bacteriophage | - | - | - |
| ALG_2_DN19013_c0_g1_i2_len1689 | ORF_2 | 224.31 | VOG22975 | 1.40E-04 | REFSEQ_carboxylesterase | Caudovirales;Siphoviridae | - | - | - |
| ALG_2_DN19410_c0_g2_i6_len1950 | ORF_1 | 183.94 | VOG12013 | 5.20E-04 | sp\|P03778\|Y06_BPT7_Protein_0.6B | Viruses | PF16752.5 | 1.50E-04 | Tubulin-specific chaperone C |
| **ALG_2_DN18226_c0_g1_i1_len1259** | **ORF_1** | **157.61** | **VOG09820** | **2.90E-04** | **REFSEQ_hypothetical_protein** | **Phycodnaviridae;Chlorovirus** | - | - | - |
| ALG_2_DN18993_c2_g2_i2_len2532 | ORF_1 | 151.17 | VOG08344 | 8.50E-08 | REFSEQ_hypothetical_protein | Bacteriophage | PF13424.6 | 3.80E-132 | Tetratricopeptide repeat |
| **ALG_3_DN34624_c0_g1_i1_len2077** | **ORF_2** | **146** | **PF02123.16** | **8.50E-05** | **Viral RNA-directed RNA-polymerase** | **RdRP-4** | - | - | - |
| **ALG_2_DN18744_c0_g1_i3_len2080** | **ORF_1** | **134.99** | **VOG10472** | **4.10E-04** | **REFSEQ_hypothetical_protein** | **Poxviridae** | - | - | - |
| ALG_2_DN19214_c2_g1_i7_len1432 | ORF_2 | 118 | VOG06927 | 6.90E-04 | REFSEQ_hypothetical_protein | Bacteriophage | - | - | - |
| **ALG_3_DN25592_c0_g1_i1_len1043** | **ORF_1** | **115** | **VOG01256** | **4.40E-04** | **sp\|Q9QU29\|ORF3_TTVB1_Uncharacterized_ORF3_protein** | **dsDNA_viruses** | - | - | - |
| ALG_2_DN18451_c0_g1_i4_len2061 | ORF_4 | 109.48 | VOG17696 | 7.10E-04 | REFSEQ_hypothetical_protein | Bacteriophage | PF16058.5 | 1.80E-17 | Mucin-like |
| ALG_2_DN18451_c0_g1_i4_len2061 | ORF_4 | 109.48 | VOG17696 | 7.10E-04 | REFSEQ_hypothetical_protein | Bacteriophage | PF16058.5 | 1.10E-07 | Mucin-like |
| **ALG_2_DN18732_c0_g2_i2_len2454** | **ORF_2** | **99.65** | **VOG09815** | **1.70E-15** | **REFSEQ_hypothetical_protein** | **Phycodnaviridae;Chlorovirus** | - | - | - |
| ALG_2_DN18957_c0_g1_i1_len1408 | ORF_2 | 97.98 | VOG02199 | 8.40E-04 | sp\|Q5UR09\|YR648_MIMIV_Uncharacterized_protein_R648 | Ortervirales | PF06156.13 | 4.70E-04 | Initiation control protein YabA |
| ALG_2_DN18957_c0_g1_i2_len1657 | ORF_2 | 89.04 | VOG02199 | 3.50E-04 | sp\|Q5UR09\|YR648_MIMIV_Uncharacterized_protein_R648 | Ortervirales | PF06156.13 | 2.20E-04 | Initiation control protein YabA |
| ALG_2_DN19250_c2_g3_i3_len2203 | ORF_1 | 71.48 | VOG23558 | 7.80E-05 | REFSEQ_hypothetical_protein | Caudovirales;Siphoviridae | - | - | - |
| ALG_2_DN19410_c0_g2_i7_len1634 | ORF_1 | 49.51 | VOG12013 | 9.90E-04 | sp\|P03778\|Y06_BPT7_Protein_0.6B | Bacteriophage | - | - | - |
| ALG_2_DN11543_c0_g1_i1_len842 | ORF_1 | 42 | VOG20356 | 3.80E-04 | REFSEQ_hypothetical_protein | Caudovirales;Myoviridae | - | - | - |
| ALG_2_DN19463_c5_g2_i1_len765 | ORF_1 | 37.7 | VOG24589 | 2.60E-05 | REFSEQ_hypothetical_protein | Caudovirales;Siphoviridae | - | - | - |
| ALG_2_DN19174_c0_g2_i18_len938 | ORF_1 | 36.39 | VOG10625 | 9.70E-04 | sp\|Q05293\|VG78_BPML5_Gene_78_protein | Bacteriophage | - | - | - |
| ALG_2_DN41289_c0_g1_i1_len750 | ORF_1 | 36 | VOG06662 | 9.50E-05 | REFSEQ_Cupin | Bacteriophage | - | - | - |
| ALG_2_DN22182_c0_g1_i1_len820 | ORF_1 | 35 | VOG02199 | 1.50E-04 | sp\|Q5UR09\|YR648_MIMIV_Uncharacterized_protein_R648 | Ortervirales | PF06156.13 | 7.20E-04 | Initiation control protein YabA |
| ALG_2_DN44027_c0_g1_i1_len815 | ORF_1 | 33.98 | VOG21678 | 3.70E-04 | REFSEQ_hypothetical_protein | Caudovirales;Myoviridae | PF08614.11 | 6.00E-05 | Autophagy protein 1(ATG16) |
| **ALG_2_DN594_c0_g2_i1_len711** | **ORF_1** | **28** | **PF17501.2** | **2.80E-04** | **Viral RNA-directed RNA polymerase** | **Viral_RdRp_C** | - | - | - |
| ALG_2_DN14271_c0_g1_i1_len772 | ORF_1 | 25.12 | VOG18617 | 2.20E-04 | REFSEQ_hypothetical_protein | Caudovirales;Siphoviridae | PF13855.6 | 1.60E-21 | Leucine rich repeat |
| ALG_2_DN18993_c2_g2_i1_len2178 | ORF_1 | 21.18 | VOG08344 | 4.70E-07 | REFSEQ_hypothetical_protein | Bacteriophage | PF13374.6 | 1.40E-126 | 1Tetratricopeptide repeat |
| ALG_2_DN44027_c0_g1_i2_len783 | ORF_1 | 19.02 | VOG21678 | 3.60E-04 | REFSEQ_hypothetical_protein | Caudovirales;Myoviridae | PF08614.11 | 7.70E-05 | Autophagy protein 1(ATG16) |
| ALG_2_DN19463_c5_g2_i6_len928 | ORF_1 | 14.34 | VOG24589 | 6.40E-05 | REFSEQ_hypothetical_protein | Caudovirales;Siphoviridae | - | - | - |
| ALG_2_DN19463_c5_g2_i4_len863 | ORF_1 | 4.36 | VOG24589 | 5.30E-05 | REFSEQ_hypothetical_protein | Caudovirales;Siphoviridae | - | - | - |

To manually assess the level of confidence of the RdRp signal detected in the HMM comparisons, the protein sequences of the three RdRp candidates were aligned to amino acid sequences retrieved from the RdRp_C, RdRp_4 and RdRp_1 PFAM profiles. The RdRp_C profile represents the C-terminal of the RdRp (Protein A) found in Alphanodaviruses. Unfortunately, this region lacks the functional motifs usually associated with the RdRp, preventing us from definitively establishing the ALG_2_DN594 contig as representing a true RdRp-encoding virus. Similarly, the ALG_3_DN34624 alignment with the viral RdRp sequences that comprise the PFAM RdRp_4 profile (PF02123) does not display a clear conservation of the crucial functional residues and motifs within the RdRp, particularly the canonical GDD motif that is GFD in ALG_3 contig (Fig S2): strikingly, the GFD motif is absent from an alignment of 4,627 viral RdRp sequences^27^. Whether this reflects a newly identified functional motif of the viral RdRp or an artefact remains to be determined, but at this point we cannot safely conclude that ALG_3_DN34624 encodes a viral RdRp. In contrast, the ALG_2_DN19089-encoded ORF shared multiple motifs with RdRp_1 profile (PF00680) (Fig S3), including motif A at positions 437-442 and GDD at positions 557-559. Because of the presence of these functional motifs, ALG_2_DN19089 can be confidently considered as a RdRp-encoding contig and will be referred to here as ‘Partiti-like adriusvirus’. Interestingly, this contig also revealed significant similarity to some eukaryotic chloroplast-associated double-stranded RNA replicons (BDRC) obtained from the green algae species, *Bryopsis cinicola*^28^ (Table 2). It is therefore likely that these BDRC dsRNA *Bryopsis*-replicons in fact represent viral RdRp sequences^20^, and we treat them as such in this study.

An additional BLASTx comparison using this divergent Partiti-like RdRp as a reference identified two other BDRC-like contigs in the *Ostreobium* sp. data set – ALG_2_DN19300 and ALG_2_DN19436 (Table 2, Fig 5). Along with the Partiti-like adriusvirus, these two additional sequences were both validated by RT-PCR (Fig S1) and are listed in the viral contig table as ‘Partiti-like lacotivirus’ and ‘Partiti-like alassinovirus’, respectively (Table 2).

### Relative abundance and prevalence of RNA viruses in the samples

Relative virus abundance varied between libraries and viruses of the same family in the *Ostreobium* sp. culture, with viral-like sequences constituting between 0.01 and 0.2% of the total non rRNA reads (Table 2, Fig 2). Each virus described was identified in only one of the cultures sequenced (Fig S1). However, inter-sample BLAST-comparisons revealed similarity between the partiti-like sequence identified in the *K. allantoideum* and *Ostreobium* samples. A non-annotated ORF from a *K. allantoideum* contig, ALG_1_DN2506, aligned with the N-terminal ORF of *Ostreobium* sp. amalga-like virus contigs that potentially encode the virus coat protein. Because of their co-occurrence in *K. allantoideum* (1.1 and 1.2 PCR, Fig S1) and their similar abundance levels, it is likely that ALG1 DN2506 and ALG 1 DN2691 (referred as ‘Amalga-like boulavirus’, Table 2) contigs are in fact part of the same genome. Unfortunately, the poor quality of RNAs and the resulting high degree of fragmentation obtained in the ALG_1 RNAseq library did not allow us to resolve this question.

Notably, a large majority of the new RNA viruses reported here come from the ulvophyte *Ostreobium* sp., although this may in part result from differences in RNA quality and sequencing rather than a true biological difference in RNA virome composition as only a limited number of non-rRNA reads were obtained for the ALG_1 and ALG_3 libraries (Fig S4). Difficulties in detecting highly divergent sequences, especially in poorly characterized and distant clades like the chlorarachniophytes, may also contribute to the different numbers of viruses observed between libraries.

### Secondary host detection

As the algal cultures analysed here were not axenic, we evaluated the potential presence of other eukaryotes in the samples using CCMetagen^29^. According to the Krona plots obtained for each library, cultivated algae were the major organism found in the samples (Fig S5). Nevertheless, 2-8% of ALG_1 and ALG_3 contigs were assigned to dinoflagellates and *Cyanophora* algae (Fig S5). Although sequences assigned to *Lingulodinium polyedrum* potentially result from GenBank mis-annotation and were likely of bacterial origin, the *Coolia malayensis* (Dinophyceae), *Amphidinium sp* (Dinophyceae) and *Cyanophora paradoxa* (Glaucophyta) associated sequences likely constitute true assignments. We therefore suspect that these additional micro-eukaryotic transcripts arose from cross-contamination with additional algae samples that were extracted and sequenced at the same time. Importantly, none of the viruses identified in the three libraries studied here could be detected in the transcriptomes of the co-processed Dinophyceae and Glaucophyta cultures, suggesting that these low-abundance contaminants are not the hosts of the viruses reported here.

### Phylogenetic analysis of the newly identified viruses

#### dsRNA viruses

##### Partiti-like viruses

Eight of the newly-described viral sequences exhibited RdRp amino acid sequence similarity with Partiti-like viruses (i.e. close to members of the *Partitiviridae*). Based on phylogenetic studies, five of the viral-like sequences are close to those from the *Amalgaviridae* (named after their mosaic status comprising both a *Partitiviridae*-like RdRp and the dicistronic and monopartite genomic organization of the *Totiviridae*^30^) and form a clade with the *Bryopsis* mitochondria-associated dsRNA (BDRM), although they share only 28-32% sequence identity (Table 2). BDRM was first described as a dsRNA associated with mitochondria in *Bryopsis cinicola* macroalgae^31^ and later classified as a virus by the International Committee on Taxonomy of Viruses (ICTV). Like *Ostreobium* and *Kraftionema*, *Bryopsis* belongs to the class Ulvophyceae, and it seems likely that all these five newly identified Amalga-like viral sequences (Fig 5) form an Ulvophyceae-infecting viral clade.

Considering the levels of RdRp pairwise identity between these sequences (Table A, Top) we assumed that each constituted a new species. Although a clear taxonomic classification needs to be established at the family or genus levels, we performed a preliminary taxonomic assessment using the PAirwise Sequence Comparison (PASC) tool from the NCBI^32^. Each of these five new BDRM-like viral genome sequences were compared with the *Amalgaviridae* full-length genomes available at NCBI. For each newly-discovered virus, the closest matches to existing *Amalgaviridae* sequences were retrieved, and the resulting pairwise identity distributions compared with those within and between *Amalgaviridae* genera (Fig 3A). While this analysis indicates that these newly identified virus sequences are not part of any existing *Amalgaviridae* genus (Fig 3A), whether they can be considered as a new genus within the *Amalgaviridae*, or a new family, is not yet clear and will require a formal taxonomic assessment by the ICTV.

**Fig 3.**
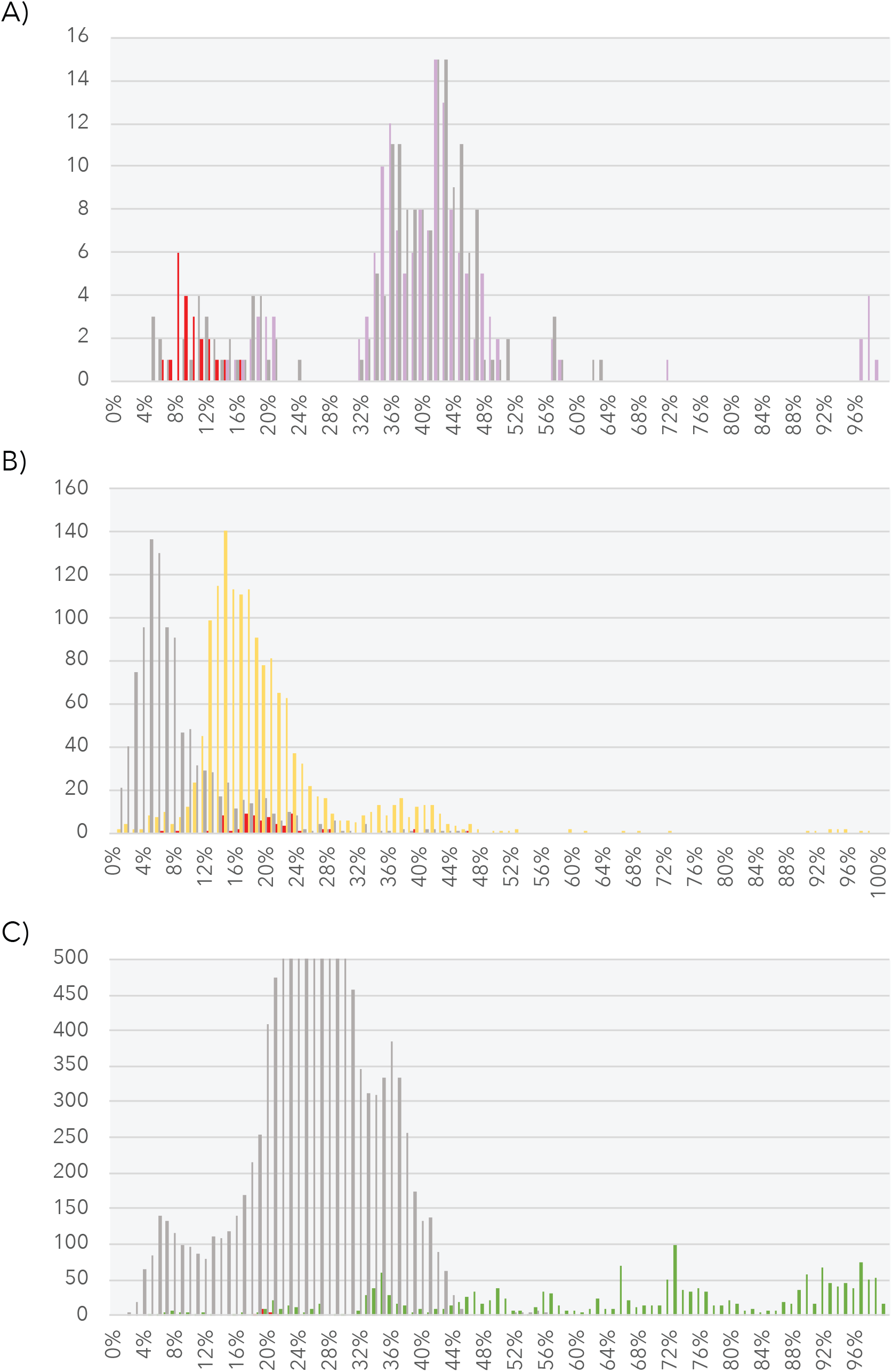
Genome pairwise identity distributions of the new algal viral sequences. The level of pairwise identity between the viruses newly identified viruses and existing members of each viral family are represented in red. (A) Inter-genus and intra-genus identity levels within the *Amalgaviridae*. Inter-genus and intra-genus identity levels are displayed in grey and purple, respectively. (B) Inter-genus and intra-genus identity levels within the *Narnaviridae*. Inter-genus and intra-genus identity levels are displayed in grey and yellow, respectively. C) Inter-genus and intra-genus identity levels within *Tombusviridae*. Inter-genus and intra-genus identity levels are displayed in grey and green, respectively.

**Fig 4.**
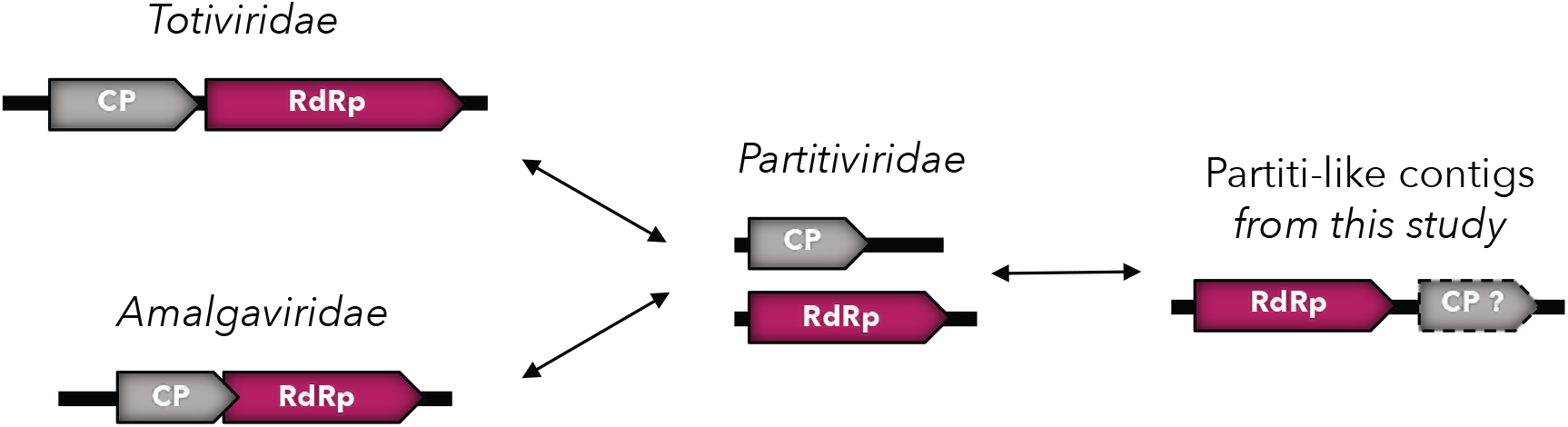
Genomic organization of the *Partitiviridae*, *Totiviridae* and *Amalgaviridae*. Possible evolutionary scenarios for the BDRC-like contigs observed in *Ostreobium* sp.

Interestingly, if these Amalga-like sequences are translated into amino acids using the protozoan mitochondrial code, they display the same genomic organization as BDRM, encoding two overlapping ORFs: the 5’ one encoding a hypothetical protein and the other coding for a replicase through a −1 ribosomal frameshift^33^ (Fig S6). However, it is unclear if these sequences should be translated through the mitochondrial genetic code or the standard cytoplasmic one code, and such sub-cellular localization still remains to be validated. It is also notable that the two closest homologs of BDRM, the Amalga-like dominovirus and Amalga-like chaucrivirus, contain the GGAUUUU ribosomal −1 frameshift motif and could plausibly be translated in this manner in the mitochondria (Fig S6).

The length and two-ORF encoding genomic structure of the BDRM-like sequences generally correspond to genomic features of the amalgaviruses (Fig 5, right). Despite a lack of detectable sequence similarity at both the sequence (BLASTx) and structural levels (Phyre2), the second ORFs predicted in these amalgavirus-like sequences are expected to encode a CP-like protein, even if the involvement of this potential CP in encapsidation remains unclear^34^. A divergent virus similar to the BDRM was also recently identified as infecting the phytopathogenic fungi *Ustilaginoidea virens*^35^.

**Fig 5.**
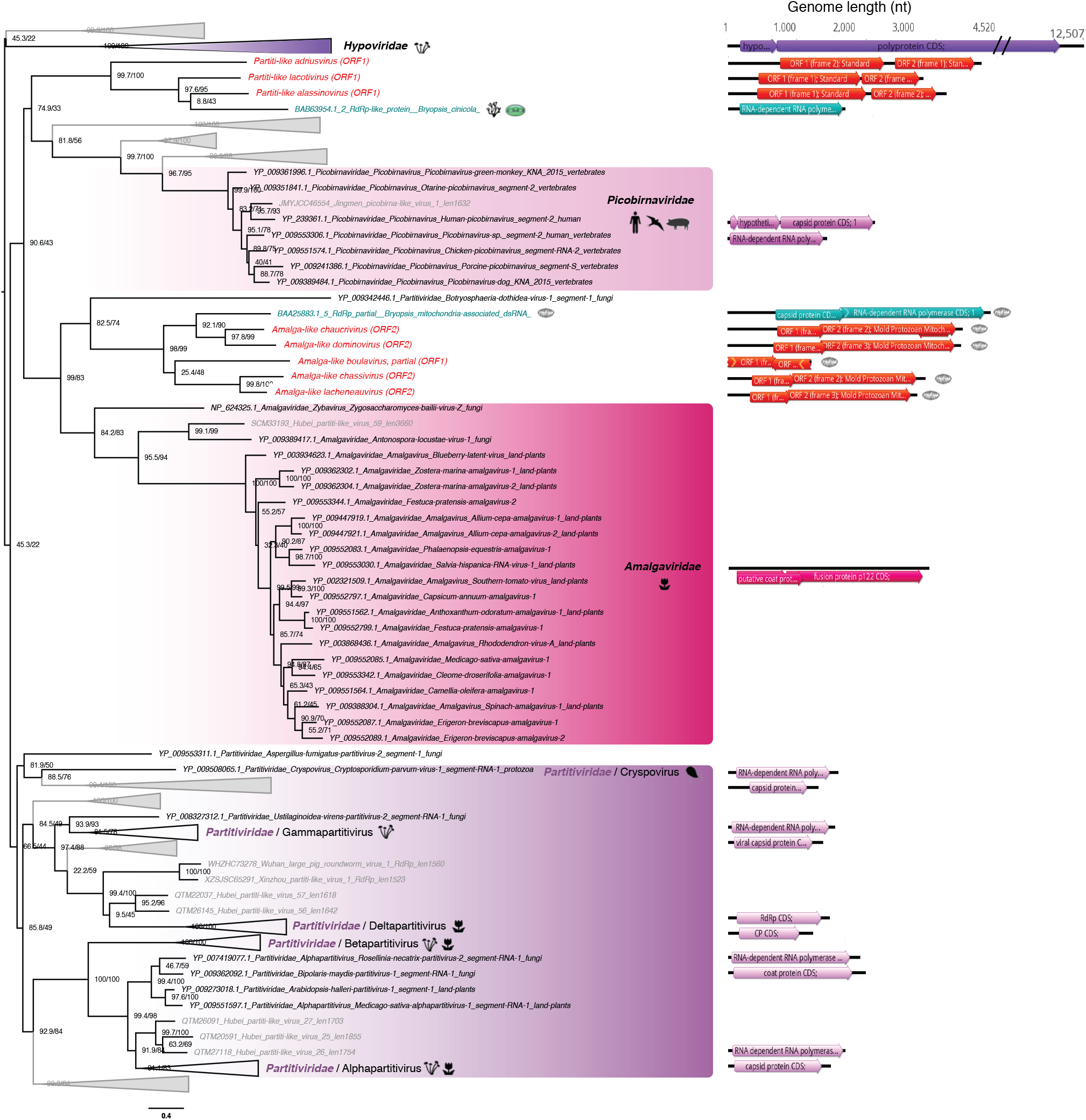
RdRp phylogeny of newly identified chlorophyte viruses among the *Amalgaviridae*, *Partitiviridae*, *Picobirnaviridae* and *Hypoviridae*. Sequences identified in this study are labelled in red. Unclassified sequences from ref.^41^ are highlighted in grey. For clarity, some families and genera have been collapsed. Right, phylogenetic tree estimated using IQ-Tree with bootstrap replicates and SH-aLTR set to 1000 (values indicated in parenthesis). Left, genomic organizations of new viruses as well as the closest identified relatives representing each family/genus as follows: Cryphonectria hypovirus 2 (*Hypoviridae*); Chicken picobirnavirus (*Picobirnaviridae*); Southern tomato virus (*Amalgaviridae*); Cryptosporidium parvum virus 1 (Cryspovirus); Discula destructiva virus 1 (*Gammapartitivirus*); Fig cryptic virus (Deltapartitivirus); Ceratocystis resinifera partitivirus (*Betapartitivirus*); White clover cryptic virus 1 (*Alphapartitivirus*). The tree is mid-pointed rooted and branch lengths are scaled according to the number of amino acid substitutions per site.

The three BDRC-like sequences identified from *Ostreobium* sp. also cluster with the Partiti-like viruses (Fig 5, Table 2) and can be classified as three different species after applying the 90% RdRp percentage identity species demarcation criteria (Table 4, Middle). Notably, the genomic organization of these BDRC-like contigs seems “inverted” compared to members of the *Totiviridae* and *Amalgaviridae*; that is, a first ORF encoding a CP protein is followed by a second that represents the RdRp. Indeed, the RdRp encoded by the Partiti-like ALG_2 contigs is close to the 5’ extremity, followed by a second ORF. This second ORF could potentially encode a CP, although functional annotation could not be achieved due to the high level of sequence divergence.

**Table 4.** RdRp-based pairwise identity of newly discovered viruses. Top: Amalga-like RdRp; Middle: BDRC-like RdRp; Bottom: Mitovirus-like RdRp.

| Amalga-like RdRp | Amalga-like chassivirus | Amalga-like lacheneauvirus | Amalga-like chaucrivirus | Amalga-like dominovirus |
| --- | --- | --- | --- | --- |
| Amalga-like boulavirus | 19 | 21 | 16 | 16 |
| Amalga-like chassivirus | - | 47 | 22 | 20 |
| Amalga-like lacheneauvirus |  | - | 22 | 21 |
| Amalga-like chaucrivirus |  |  | - | 36 |

| BDRM-like RdRp | Partiti-like alassinivirus | Partiti-like lacotivirus |
| --- | --- | --- |
| Partiti-like adriusvirus | 20 | 20 |
| Partiti-like alassinivirus | - | 32 |

| Mitovirus-like RdRp | Mito-like<br>picolinus<br>virus | Mito-like<br>babylonus<br>virus | Mito-like<br>albercanus<br>virus | Mito-like<br>laruketanus virus | Mito-like<br>boobanusbagrillonus<br>virus | Mito-like<br>bionusvirus |
| --- | --- | --- | --- | --- | --- | --- |
| Mito-like spartanusvirus | 46 | 19 | 21 | 21 | 20 | 15 |
| Mito-like picolinusvirus | - | 17 | 20 | 19 | 19 | 13 |
| Mito-like babylonusvirus |  | - | 22 | 22 | 20 | 14 |
| Mito-like albercanusvirus |  |  | - | 26 | 25 | 16 |
| Mito-like laruketanusvirus |  |  |  | - | 37 | 14 |
| Mito-like<br>boobanusbagrillonusvirus |  |  |  |  | - | 12 |

##### Structural-based homology analysis

The remaining hits from the RdRp-profile analysis – ALG_2_DN594_c0_g2_i1_len711 and ALG_3_DN34624_c0_g1_i1_len2077 – were used in a Phyre2 analysis. However, this revealed no confident identification of viral RdRp signal (i.e. the confidence levels of models were < 90%).

Despite the great diversity of host and genomic organizations, the detectable sequence similarity observed between *Partitiviridae* and *Amalgaviridae* strongly suggests that they share common ancestry^34^. As such, it is important to determine whether amalgaviruses are restricted to plant hosts^36–38^, and hence differ from the *Partitiviridae* that have been characterized in many divergent eukaryotic hosts such as fungi, plants and protists^39,40^.

#### ssRNA(+) viruses

##### Mitovirus-like viruses

Seven viral sequences from *Ostreobium* sp. clustered in the *Narnaviridae*, forming a clade within the genus *Mitovirus* (Fig 6). Their single ORF encodes an RdRp, and a genome length of ~3000 nt is typical of the *Narnaviridae* (Fig 6, right). These viral sequences could constitute a new clade of protist-associated mitoviruses potentially restricted to green microalgae. Indeed the closest relative identified here, Shahe Narna-like virus 6, was isolated from freshwater small planktonic crustaceans (*Daphnia magna*, *Daphnia carinata* and *Moina macrocopa*) belonging to the order Cladocera^41^ (commonly referred to as water fleas). These animals feed on microalgae and it is therefore possible that Shahe Narna-like virus 6 in fact infects algae rather than arthropods, in a similar manner to other members of the *Narnaviridae*. It is therefore possible that these RNA viruses could be major constituents in habitats occupied by protists and by consequence in larger predatory invertebrates.

**Fig 6.**
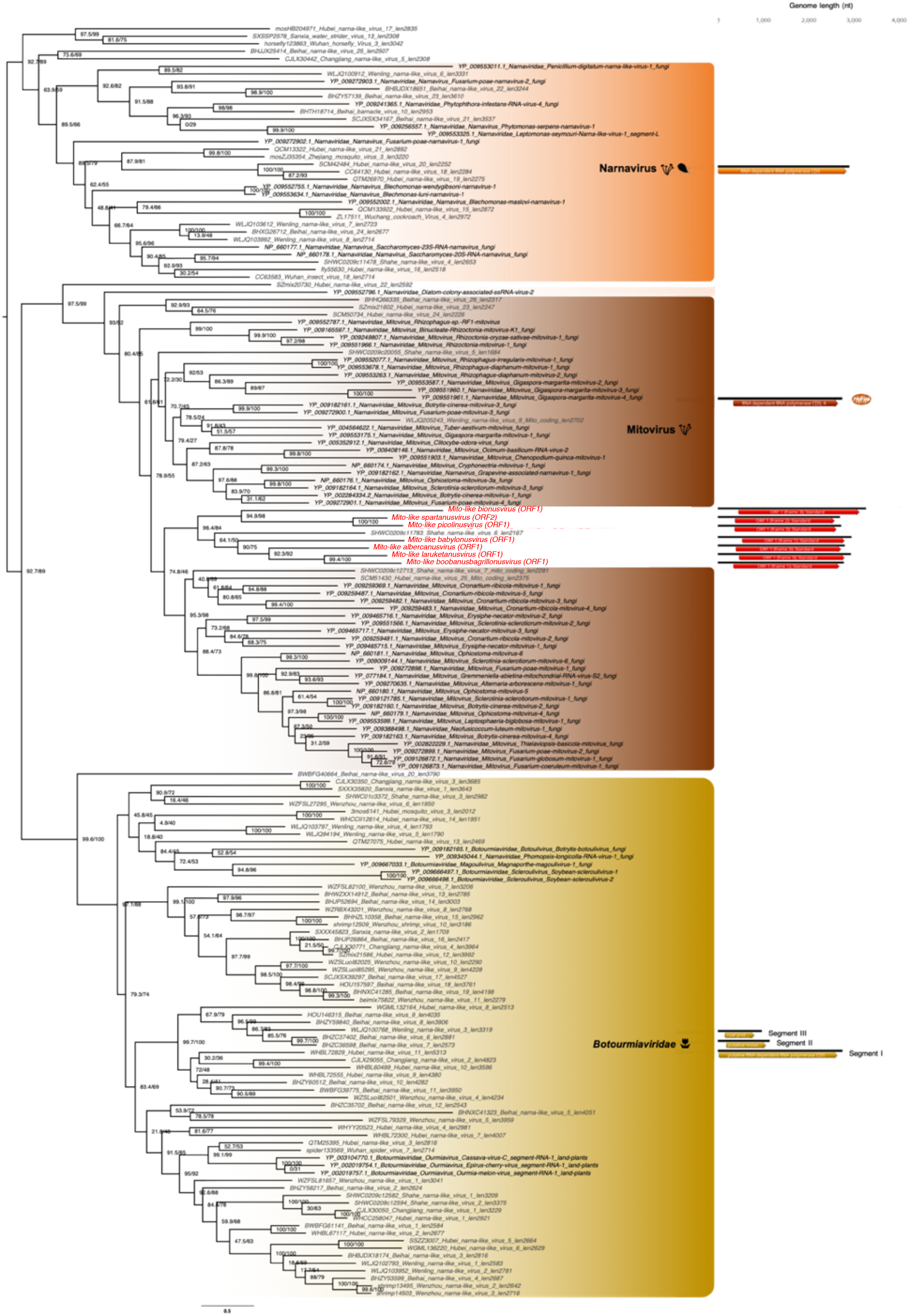
Phylogeny of the *Narnaviridae*-*Botourmiaviridae* group based on the RdRp. Newly discovered viruses from *Ostreobium* sp. are highlighted in red. RdRp sequences from unassigned RNA virus retrieved from ref.^41^ are marked in grey. Right, phylogenetic tree estimated using IQ-Tree with bootstrap replicates and SH-aLTR set to 1000 (values indicated in parenthesis). Left, genomic organization of both viral genomes identified in this study (red) and representatives of each major family and genus used in the phylogeny (Cassava virus C – *Botourmiaviridae*; Saccharomyces 23S RNA narnavirus; Chenopodium quinoa mitovirus 1). Annotations of Cassava virus C coding sequences: RdRp (Segment I); Putative movement protein (Segment II); Coat protein (Segment III). Branch lengths are scaled according to the number of amino acid substitutions per site.

Based on the 50% RdRp sequence identity used as a species demarcation criterion in the *Narnaviridae* (ICTV report 2009), we identified seven new mito-like viral species (Table 4, bottom). To place of these new species among the *Narnaviridae*, we performed a PASC analysis using full-length genomes from the seven new viral species. This revealed that the identity levels of the new sequences fell in the range expected of intra-genus diversity (Fig 3B). We therefore propose the existence of a new sub-clade of mitoviruses, including these seven new species as well as the Shahe Narna-like virus 6 previously described^41^ (Fig 6). Whether this proposed clade is associated with mitochondria is currently unclear, and predicted ORFs were obtained using both standard and mitochondrial genetic codes.

#### Tombusviridae-like viruses

One sequence from *Ostreobium* sp., the Tombus-like chagrupourvirus, exhibited similarity to members of the *Tombusviridae* family of ssRNA(+) viruses (Fig 7), grouping with viruses previously identified as infecting plant or plant pathogenic fungi. This again illustrates the likely shared evolutionary history between green algae and land plants, and that horizontal virus transfers can occur between plants and their pathogenic fungi. Of note is that the closest relative of Tombus-like chagrupourvirus documented to date, Hubei-Tombus like virus 12, was isolated from freshwater animals (mollusca *Nodularia douglasiae)*^41^. According to the lack of distinguishable animal-related contigs in the *Ostreobium* sp. sample (Fig S5), this Hubei-Tombus like virus 12 may, together with the newly tombus-like sequence discovered here, constitute a new clade of green algae-infecting viruses.

**Fig 7.**
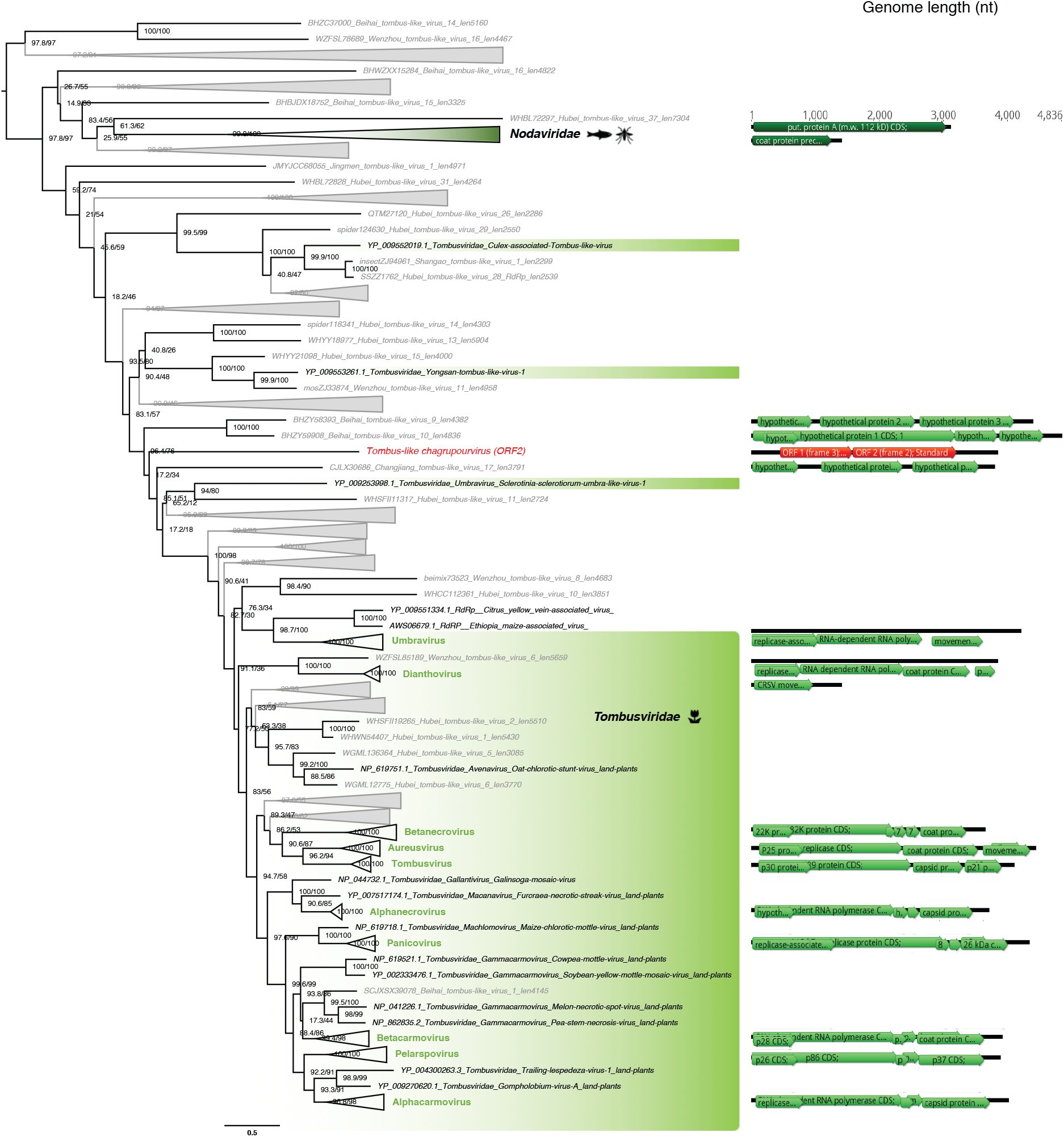
Phylogeny of the *Tombusviridae* RdRp. The tombus-like sequence identified in this study is labelled in red. Unclassified sequences from ref.^41^ are highlighted in grey. For clarity, some families and genera have been collapsed. Right, phylogenetic tree estimated using IQ-Tree with bootstrap replicates and SH-aLTR set to 1000 (values indicated in parenthesis). Left, genomic organizations of new viruses as well as their closest homologs and representatives from each family/genus as follows: Black beetle virus (*Nodaviridae*); Carrot mottle virus (genus *Dianthovirus*); Carnation ringspot virus (Dianthovirus); Beet black scorch virus (*Betanecrovirus*); Cucumber leaf spot virus (*Aureusvirus*); Maize necrotic streak virus (*Tombusvirus*); Olive latent virus 1 (*Alphanecrovirus*); Panicum mosaic virus (*Panicovirus*); Hibiscus chlorotic ringspot virus (*Betacarmovirus*); Clematis chlorotic mottle virus (*Pelarspovirus*); Carnation mottle virus (*Alphacarmovirus*). The tree is mid-pointed rooted and branch lengths are scaled according to the number of amino acid substitutions per site.

The 3.8kb genome length of the Tombus-like chagrupourvirus is similar to those commonly observed in *Tombusviridae* and their relatives (Fig 7, right), suggesting that it comprises a full-length genome for this virus. This putative full-length genome sequence was compared to *Tombusviridae* reference genomic sequences using PASC to assess its taxonomic position (Fig 3C). Accordingly, the *Ostreobium*-associated tombus-like sequence could constitute a new *Tombusviridae* genus.

#### Virgaviridae-like viruses

One sequence identified in *Chlorarachnion reptans* displayed detectable sequence similarity to the *Virgaviridae*-like RdRp supergroup (Fig 8). The family *Virgaviridae c*omprise ssRNA(+) viruses traditionally associated with plants and display diverse genomic organizations. The short length of the Virga-like bellevillovirus associated with chlorarachniophytes indicates that this sequence likely reflects a partial genome sequence only. Moreover, the multi-segment structure of the closest relatives suggests that the partial genome recovered in ALG_3 could also contain additional segments not yet identified. Although further host confirmation is required, this newly described RNA virus-like sequence would constitute the first algae virus from the Hepe-Virga group.

**Fig 8.**
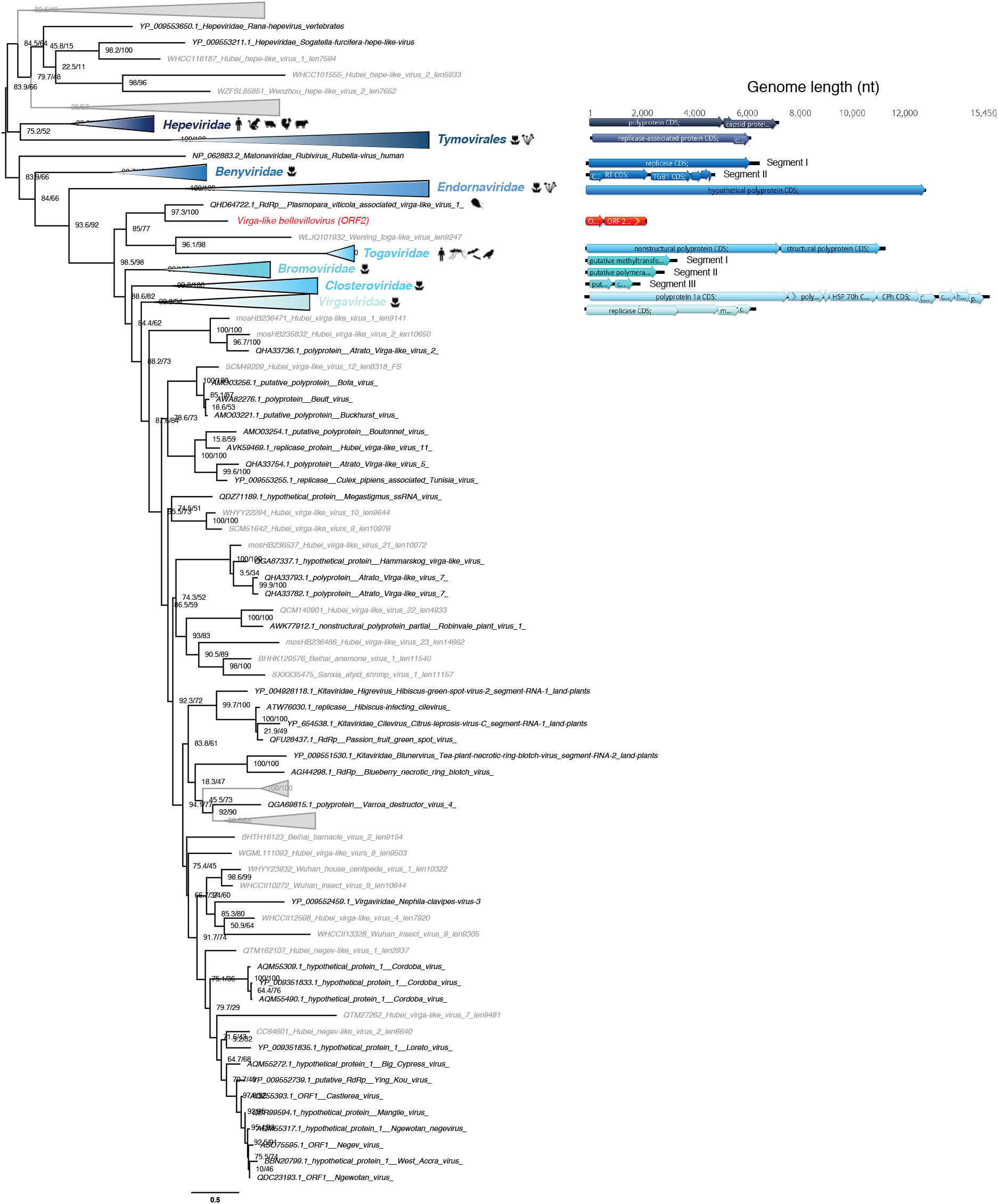
Phylogeny of the Hepe-Virga group RdRp. The hepe-virga-like sequence identified in this study is labelled in red. Unclassified sequences from ref.^41^ are highlighted in grey. For clarity, some families or genera have been. Right, RdRp-based phylogenetic tree obtained using IQ-tree with bootstrap replicates and SH-aLTR set to 1000 (resulting values are indicated in parenthesis). Left, genomic organizations of new viruses as well as closest homologs and representative from each family/genus as follows: Orthohepevirus A (*Hepeviridae*); Poinsettia mosaic virus (order *Tymovirales*); Wheat stripe mosaic virus (*Benyviridae*); Diatom colony associated dsRNA virus 15 (*Endornaviridae*); Cabassou virus (*Togaviridae*); Apple mosaic virus (*Bromoviridae*); Mint virus 1 (*Closteroviridae*); Cucumber mottle virus (*Virgaviridae*). The tree is mid-pointed rooted and branch lengths are scaled according to the number of amino acid substitutions per site.

#### ssRNA(-) virus

##### Bunyavirales-like viruses

A partial viral genome, the Bunya-like bridouvirus, encoding a RdRp-like signal was identified in the *Chlorarachnion reptans* sample, although this sequence is highly divergent and cannot be formally assigned to any previously described viral family. Strikingly, however, the sequence clusters with a Bunya-Arena like virus previous identified in diverse Freshwater small planktonic crustaceans (*Daphnia magna* (N/A), *Daphnia carinata* (N/A) and *Moina macrocopa* (N/A))^41^ that typically feed on algae. Our phylogenetic analysis places this sequence within the diversity of the order *Bunyavirales* (Fig 9). In addition to the freshwater organism-associated viruses identified in Shi et al. 2016 (ref.^41^), this contig clusters with several bunya-like unclassified negative-strand viruses isolated from the fungi *Cladosporium cladosporioides* and the oomycete *Plasmopara viticola* (Fig 9) that are both plant pathogens. The multi-segment structure of the closest classified family, the *Phenuiviridae*, suggests that additional segments associated with the partial Bunya-like bridouvirus genome may exist in *C. reptans*.

**Fig 9.**
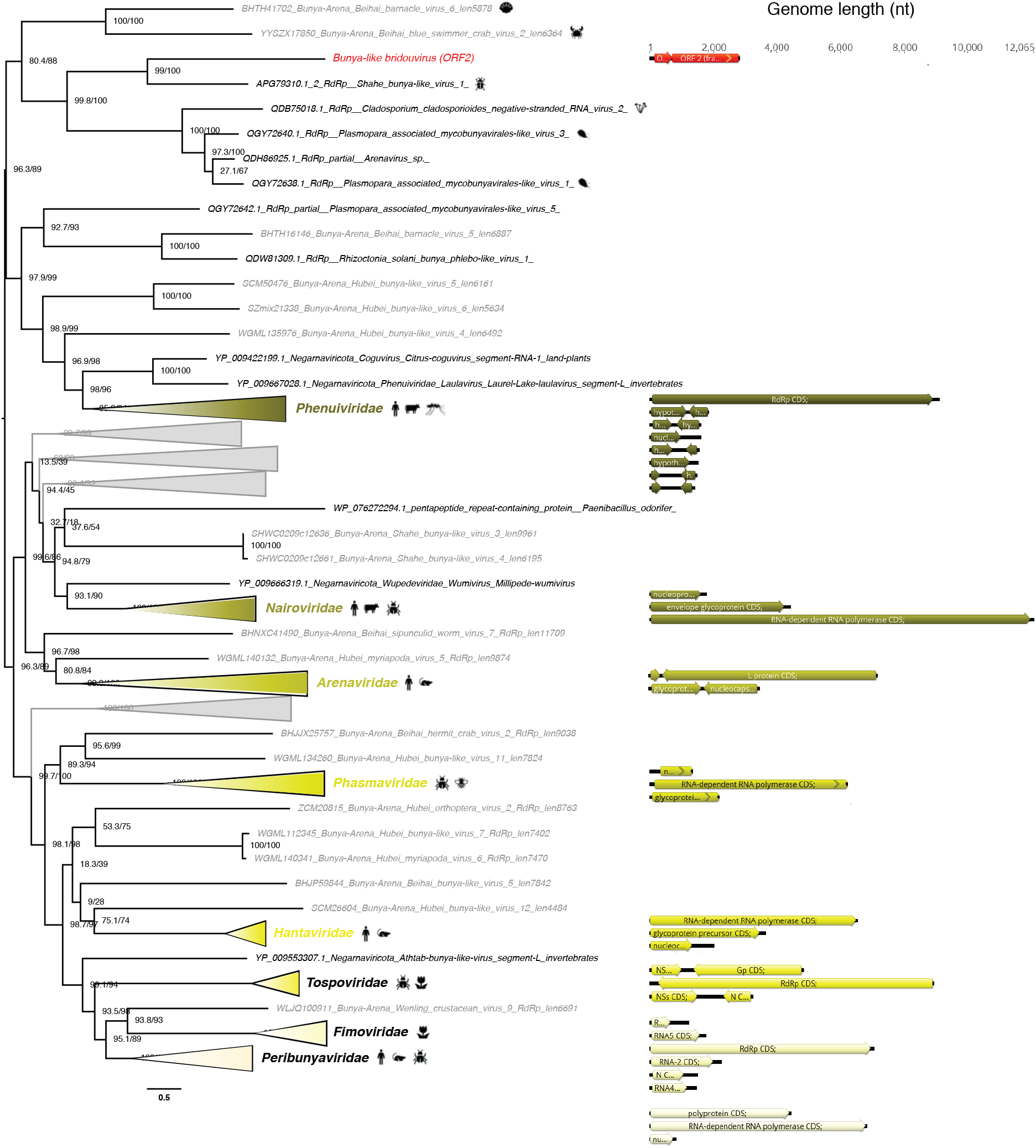
Phylogeny of the *Bunyavirales* RdRp. The bunya-like sequence identified in this study is labelled in red. Unclassified sequences from ref.^41^ are highlighted in grey. For clarity, some families and genera have been collapsed. Right, phylogenetic tree estimated using IQ-Tree with bootstrap replicates and SH-aLTR set to 1000 (values indicated in parenthesis). Left, genomic organizations of new viruses as well as their closest homologs and representatives from each family/genus as follows: Melon chlorotic spot virus (*Phenuiviridae*); Yogue virus (*Nairoviridae*); Latino mammarenavirus (*Arenaviridae*); Seattle orthophasmavirus (*Phasmaviridae*); Melon yellow spot virus (*Tospoviridae*); Fig mosaic emaravirus (*Fimoviridae*); Tataguine orthobunyavirus (*Peribunyaviridae*). The tree is mid-pointed rooted and branch lengths are scaled according to the number of amino acid substitutions per site.

## Discussion

The central aim of this study was to better characterise the RNA virus diversity in two major algal lineages, the chlorarachniophytes and the ulvophytes, for which no RNA viruses had previously been reported. Our investigation of the RNA virus diversity in six microalgae species led to the identification of 18 new divergent RNA viruses, although they exhibited clear homology with five established viral families. This clearly supports the idea that are an important reservoir of untapped RNA virus diversity and that the apparent domination of DNA viruses likely originates, at least partially, from sampling biases. The notion of a potentially large dark matter of algal viruses is further reinforced by the high proportion of unassigned contigs obtained, that likely contain a non-neglectable proportion of highly divergent viral reads.

### RNA virome similarities between green algae and land plants

Among the 18 novel RNA virus species described here, seven of those detected in the green algae *Ostreobium* sp. and *K. allantoideum* were seemingly related to the *Tombusviridae* and *Amalgaviridae* families of plant RNA viruses. Such similarities in RNA virome between green algae and land plants are consistent with previous analyses based on the Plant Genome project transcriptomic data that identified Partitivirus-like signals in Chlorophyte algae^20^. However, the very limited sequence similarity among these viruses strongly suggests an ancient divergence among these viral families, perhaps even before the chlorophyte-streptophyte split some 850-1,100 million years ago (Ma)^25^. The close link between land plant and green algae RNA virosphere is reinforced by the recent observation that plant viruses are able to infect non-vascular plants such as mosses and algae^42,43^.

### Divergent fungal mitoviruses in Ostreobium sp

The presence of a potential new clade of protist-associated mitoviruses is of importance as mitoviruses have traditionally been viewed as restricted to fungal hosts and were only very recently identified in plants^44^. Similar to virus transfer between fungi and land plants ^5^, it is possible that the symbiosis and co-evolution between green algae and fungi^46^ explains the close phylogenetic relationships in their virome, perhaps facilitating horizontal gene transfer events. Indeed, coral holobionts are the place for frequent interactions between endolithic algae, such as *Ostreobium* sp., and fungi^47–49^. While we cannot formally identify host associations from our RNA-seq data alone, no fungal-assigned contigs were detected in our data, arguing against these being fungal viruses. Considering the high levels of sequence divergence between our viral sequences and those associated with fungi and within the clade formed by *Ostreobium* sp.-associated viruses, it seems likely that any such horizontal gene transfer events are not recent and may have occurred in Ulvophyceae or even Chlorophyta ancestors. It will be of considerable interest to examine this new mitovirus-clade across a larger set of green microalgae species. Further characterisation of such green algae-infecting mitovirus diversity and their lifecycle could be pivotal for our understanding of the *Narnaviridae* and their evolution in eukaryotic cells. For example, it is of interest to determine whether this putative mitochondrial subcellular location is the result of an escape from the cytoplasmic dsRNA silencing (as suggested for newly characterized plant mitoviruses^50^) or if these viruses are relics of the eukaryotic endosymbiosis event, particularly as mitoviruses have bacterial counterparts, the *Leviviridae*^27^. More broadly, these newly-reported mitovirus-like sequences further illustrate the enormous diversity of hosts infected by *Narnaviridae*, including such eukaryotic microorganisms as Apicomplexa, Excavates and Oomycetes hosts^51–54^.

### Detection of plant viruses in the chlorarachniophytes

The apparent similarity between a Rhizarian (*C. reptans*) associated viral sequence and land-plant infecting viruses was also striking. The Rhizaria and Archaeplastida are assumed to have diverged before the cyanobacteria primary endosymbiosis event, ca. 1.5 billion years ago^55,56^. Thus, a detectable sequence similarity between Rhizaria-associated viruses and those infecting land plants cannot be reasonably attributed to such ancient evolutionary events. Instead, the presence of such land plant-like viruses in Chlorarachniophyte would more likely reflect more recent acquisition of these viruses in chlorarachniophytes through either horizontal transfer by common vector/symbiont/parasite ancestors or secondary endosymbiosis (eukaryote-to-eukaryote) events. Indeed, a secondary endosymbiosis event of a green alga in the core Chlorophyta, possibly related to Bryopsidales, led to the origin of the plastid of chlorarachniophytes between 578-318 Ma^21^. Whether this virus (i) constitutes a relic of viruses that infected this engulfed green algal endosymbiont, (ii) is part of the chlorarachniophyte cytoplasm or still associated with the periplastidial compartment (i.e. remnant cytoplasm of the endosymbiont), or (iii) interacts with the nucleomorph (remnants of the green algal endosymbiont nucleus) are key questions in the evolution of eukaryotic RNA viruses. While our data cannot provide answers, it will be of major interest to extend these analyses to euglenophytes and the dinoflagellate genus *Lepidodinium* where distinct secondary endosymbiosis with green algae have also occurred, as well as to cryptophytes that also contain the remnant nucleus of its red algae endosymbiont^57,58^.

### First report of a negative-sense RNA virus in algae

Our detection of Bunyavirales-like sequence in *C. reptans* is of importance as it would constitute the first evidence of a negative-sense RNA virus in a microalgae and only the second discovered in eukaryotic microorganisms. Considering the extensive host range attributed to *Bunyavirales* (from land plants to invertebrates, vertebrates and humans), their association with chlorarachniophyte hosts is plausible. However, considering the poor level of abundance in *C. reptans* sample, additional work is clearly needed to retrieve full-genome information and to confirm the association of such bunya-like viruses with chlorarachniophytes.

One of the greatest challenges in viral meta-transcriptomics is our capacity to detect distant homologies, especially in rapidly evolving RNA viruses. As a first attempt to retrieve such ephemeral evolutionary signals, we scanned orphan contigs using RdRp protein structure information in addition to the standard primary amino acid sequence. Notably, both RdRp profile and 3D protein structure comparison led to the identification of highly divergent RNA virus candidates, although these remain difficult to annotate. Efforts to better describe the repertoire of sequence and structure of viral RdRps are central to unveiling the RNA virosphere in overlooked eukaryotic organisms.

## MATERIAL AND METHODS

### Algal cultures

Algal strains were isolated from marine sand (*Microrhizoidea pickettheapsiorum*^59^; *Kraftionema allantoideum*^60^; *Chlorarachnion reptans* and *Lotharella* sp. from the Wye River, Victoria, Australia) or coral skeletons (*Ostreobium* sp. HV05007, Kavieng, Papua New Guinea), or obtained from the NIVA Culture Collection of Algae (*Dolichomastix tenuilepis*, SCCPA strain K-0587). Cultures were maintained in K-enriched seawater medium (transferring every other week) at either 26°C (*Ostreobium*) or 16°C (all others) under cool white LED lights at 1 Photosynthetic active radiation (PAR) (*Ostreobium*) or 15 PAR (all others). Cultures were pelleted by centrifugation in falcon tubes and stored in RNAlater at – 80°C until RNA extraction.

### Total RNA extractions

For total RNA extractions, RNAlater was removed by low centrifugation, algal cells were disrupted using thaw/freezing cycles and bead beating (0.1-0.5mm), and total RNA was further extracted using the Qiagen® RNeasy Plant mini kit following the manufacturer’s instructions.

For the initial using meta-transcriptomic screening, RNAs were pooled into three groups: (i) the chlorophyta *Dolichomastix tenuilepis* and *Microrhizoidea pickettheapsiorum* (Mamiellophyceae) and *Kraftionema allantoideum* (Ulvophyceae) were pooled into metatranscriptome ‘ALG_1’; (ii) the ulvophyte *Ostreobium* sp. comprised ‘ALG_2’; and (iii) the two chlorarachniophytes *Chlorarachnion reptans* and *Lotharella* sp. were pooled into ‘ALG_3’ (Table 1).

### Total RNA Sequencing

RNA quality was assessed and TruSeq stranded libraries were synthetized by the Australian Genome Research Facility (AGRF), using either (i) TruSeq stranded with a eukaryotic rRNA depletion step (RiboZero Gold kit, Illumina) for ALG_2, or (ii) the SMARTer Stranded Total RNA-Seq Kit v2 – Pico Input Mammalian libraries (Takara Bio USA) for ALG_1 and ALG_3 (Table S1), due to the low amount of input RNAs in these libraries. The resulting libraries were sequenced on an Illumina HiSeq2500 (paired-end, 100bp) at the AGRF. Library descriptions and RNA-seq statistics are summarized in Table S1.

### In silico processing of meta-transcriptomic data

#### Read depletion and contig assembly

The RNA-seq data were first subjected to low-quality read and Illumina adapter filtering using the Trimmomatic program^61^. Ribosomal RNA was depleted using the SILVA database v32^62^, which removed between 86 and 94% of the total unfiltered reads (Fig S1A/2). Read-depleted libraries were then *de novo* assembled using the Trinity program^63^ and contigs shorter than 200nt were removed (the average length of contig assembly is shown in Fig S1B/2). Contig abundances were calculated from the RNA-seq data using the Expectation-Maximization (RSEM) software^64^.

#### RNA virus detection using BLASTx and BLASTn

The similarity of contigs to the current NCBI nucleotide (nt) and protein (nr) databases was determined using the BLASTn and Diamond BLASTx programs^65^, respectively, employing 1e-10 and 1e-05 as e-value cut-offs. RNA virus-like sequences were identified using BLASTx against all RdRp protein sequences available on NCBI/GenBank. False-positive signals for RNA viruses were removed by BLASTing RdRp-like sequences against the nr database and discarding sequences displaying a non-viral sequence as the best hit.

#### RNA virus profile-based homology detection

To detect especially diverse RdRp-based sequences, orphan contigs (i.e. those with no match in either the nr and nt databases) were compared to the PFAM RdRp-protein profiles ‘MitoVir_RdRp’ (PF05919), ‘Birna_RdRp’ (PF04197), ‘Viral_RdRp_C’ (PF17501), ‘RdRP_1’ (PF00680), ‘RdRP_2’ (PF00978), ‘RdRP_3’ (PF00998) and ‘RdRP_4’ (PF02123), as well as to the entire VOG profile database (http://vogdb.org)^66^ using the HMMer3 program^67^. To check for false-positive signals, these orphan sequences were submitted to the entire PFAM database.

#### 3D protein structure prediction of RdRp-like contigs

To infer a structural model for the distant RdRp signals detected using profiles, sequences displaying a RdRp-like signal were subjected to the normal mode search of the Protein Homology/analogY Recognition Engine v 2.0 (Phyre2) web portal^68^. Briefly, this program first compares the submitted amino acid sequence to a curated non-redundant nr20 data set using HHblits^69^. It then converts the conserved secondary structure information as a query against known 3D-structures using HHsearch^70^. A final structural modelling step based on identified structural homologies is then performed as described previously^68^.

#### RNA virus sequence analysis and annotation

Total non-rRNA reads were mapped onto RNA virus-like contigs using Bowtie2 and heterogeneous coverage and potential mis-assemblies were manually resolved using Geneious v11.1.4^71^.

Open reading frames (ORFs) were first predicted using getORF from EMBOSS, in which ORFs were defined as regions that are free of stop codons, although partial sequences (i.e. missing start or stop codons) were retained for analysis. Protein domains were then annotated using the InterProscan software package from EMBL-EBI, comprising ProSite, PFAM, TIGR, SuperFamily, and CDD among others (https://github.com/ebi-pf-team/interproscan).

#### Revealing host-virus associations

A challenge faced by all metagenomic studies is confidently assigning each viral sequence to a particular host in a given sample. We used algal cultures to minimize the number of potential additional cellular hosts. These cultures were, however, non-axenic, with mainly bacteria present. To evaluate the possibility of additional microeukaryotic cells in the sample, we obtained taxonomic identification for contigs in the metatranscriptome by aligning them to the NCBI nt database using the KMA aligner and the CCMetagen program^29,72^. Contigs matching an entry in the nt database were displayed as Krona plots and classified based on their taxonomy (utilising high taxonomic levels for clarity).

#### Phylogenetic analysis

RdRp amino acid sequences were aligned using the L-INS-I algorithm in the MAFFT program^73^ and trimmed with TrimAI (automated mode). Maximum likelihood phylogenetic trees were then estimated using IQ-TREE, employing ModelFinder to obtain the best-fit model of amino acid substitution in each case, with nodal support assessed using 1000 bootstrap replicates and 1000 replicates of SH-like approximate likelihood ratio test (SH-aLRT)^74,75^. For each tree, reference genomes and corresponding RdRp sequences were retrieved from the NCBI Viral genome resource (https://www.ncbi.nlm.nih.gov/genome/viruses/). To depict the evolutionary relationships of the newly discovered viruses as meaningfully as possible, the closest unclassified BLASTx homologs were used in the phylogenetic analysis. This resulted in alignments of 237, 76, 198, 558 and 325 RdRp protein sequences for Amalga-like, Mito-like, Tombus-like, Virga-like and Bunya-like virus groups, respectively.

### RT-PCR validation

Viral contigs were validated experimentally and associated to individual algal sample using Reverse Transcription (SuperScript™ IV reverse transcriptase – Invitrogen™) followed by PCR (Platinum™ SuperFi™ DNA polymerase – Invitrogen™), with specific primer sets for each contig. The *rbc*L and *tuf*A marker genes were used as PCR positive controls using sets of primers designed in ref.^76^. All primers used in this study are described in Table S2.

## ACKNOWLEDGMENTS

We acknowledge the SIH Bioinformatics Hub and the University of Sydney’s high-performance computing cluster Artemis for providing the computing resources used for this study. We also thank Fabien Burki for providing us source files to build Figure 1.

## SUPPLEMENTARY FIGURE LEGENDS

**Fig S1. Gel electrophoresis of RT-PCR to validate the putative new RNA viruses in each sample used for library construction and RNA-seq.** RT-PCR validation in (A) ALG_1; (B) ALG_2; (C) ALG_3. rbcL: ribulose-bisphosphate carboxylase gene, tufA: elongation factor Tu, both used as positive PCR-controls. K.a: *Kraftionema allantoideum*; M.p: *Microrhizoidea pickettheapsiorum*; D.t :*Dolichomastix tenuilepsis*.

**Fig S2. MAFFT-alignment of the ALG_3_DN34624-encoded ORF with the viral RdRp sequences that compose PFAM RdRp_4 profile.**

**Fig S3. MAFFT-alignment of the ALG_2_DN19089-encoded ORF with the PFAM RdRp_1 profile entries.**

**Fig S4. rRNA depletion and contig assembly results.** (A) Read counts before and after trimming and SortmeRNA rRNA depletion. (B) Trinity assembly quality expressed as the average length of contig assembled for each library.

**Fig S5. Relative abundance of taxonomically-assigned contigs.** Krona graphs obtained using KMA and CCMetagen methods for (A) ALG_1, (B) ALG_2 and (C) ALG_3 libraries. The ORFans contigs (i.e. those without any match in nt database) are not shown here. Each circle represents a taxonomic level which is grouped and coloured based on their taxonomic classification at lower ranks.

**Fig S6. Genomic organization of the ALG_2 Partiti-like sequences and Bryopsis mitochondria-associated dsRNA.** ORFs are predicted using the mold protozoan mitochondrial genetic code (translation table 4). Ribosomal −1 frameshifting motifs “GGAUUUU” are indicated in red boxes.

## SUPPLEMENTARY TABLES

**Table S1.** RNAseq library preparation and sequencing results.

| Library ID | Library prep protocol | Nb paired-end reads | Data yield (bp) |
| --- | --- | --- | --- |
| **ALG_1** | SMARTer Stranded Total RNA-Seq Kit v2 Pico Input Mammalian | 21,014,207 | 5.30 Gb |
| **ALG_2** | TruSeq stranded/RiboZero gold | 25,963,002 | 6.54 Gb |
| **ALG_3** | SMARTer Stranded Total RNA-Seq Kit v2 Pico Input Mammalian | 18,448,371 | 4.65 Gb |

**Table S2.**
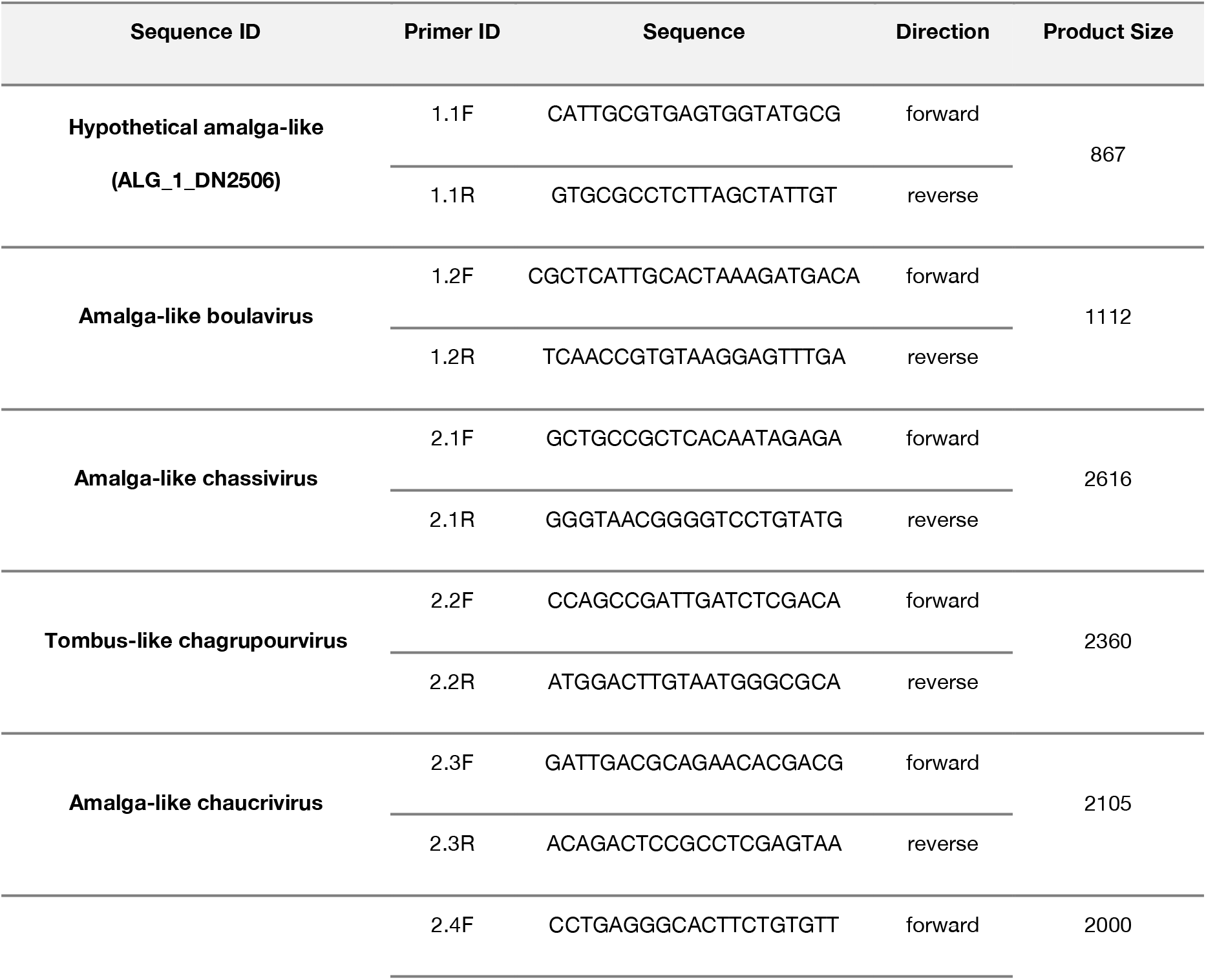
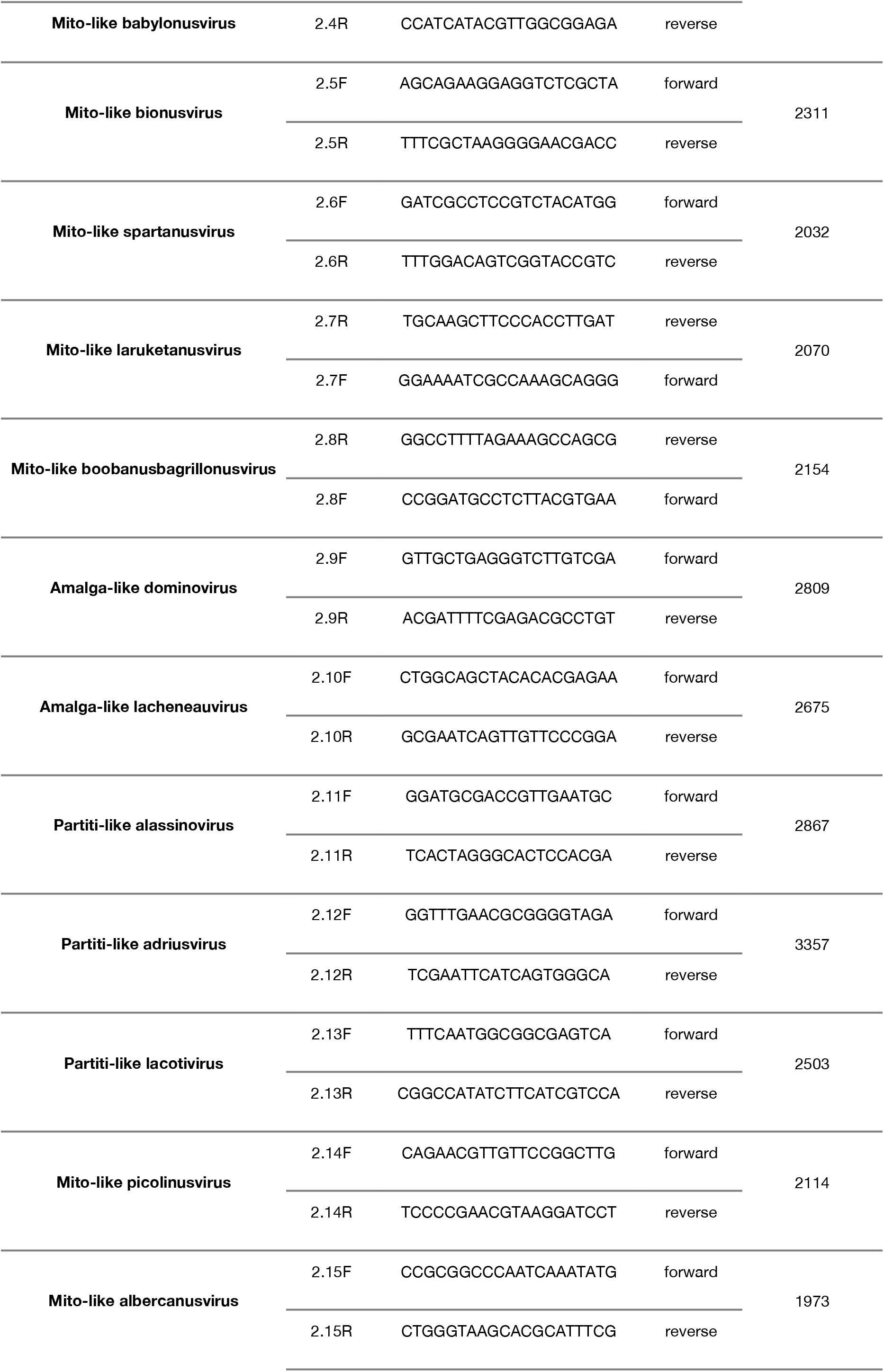
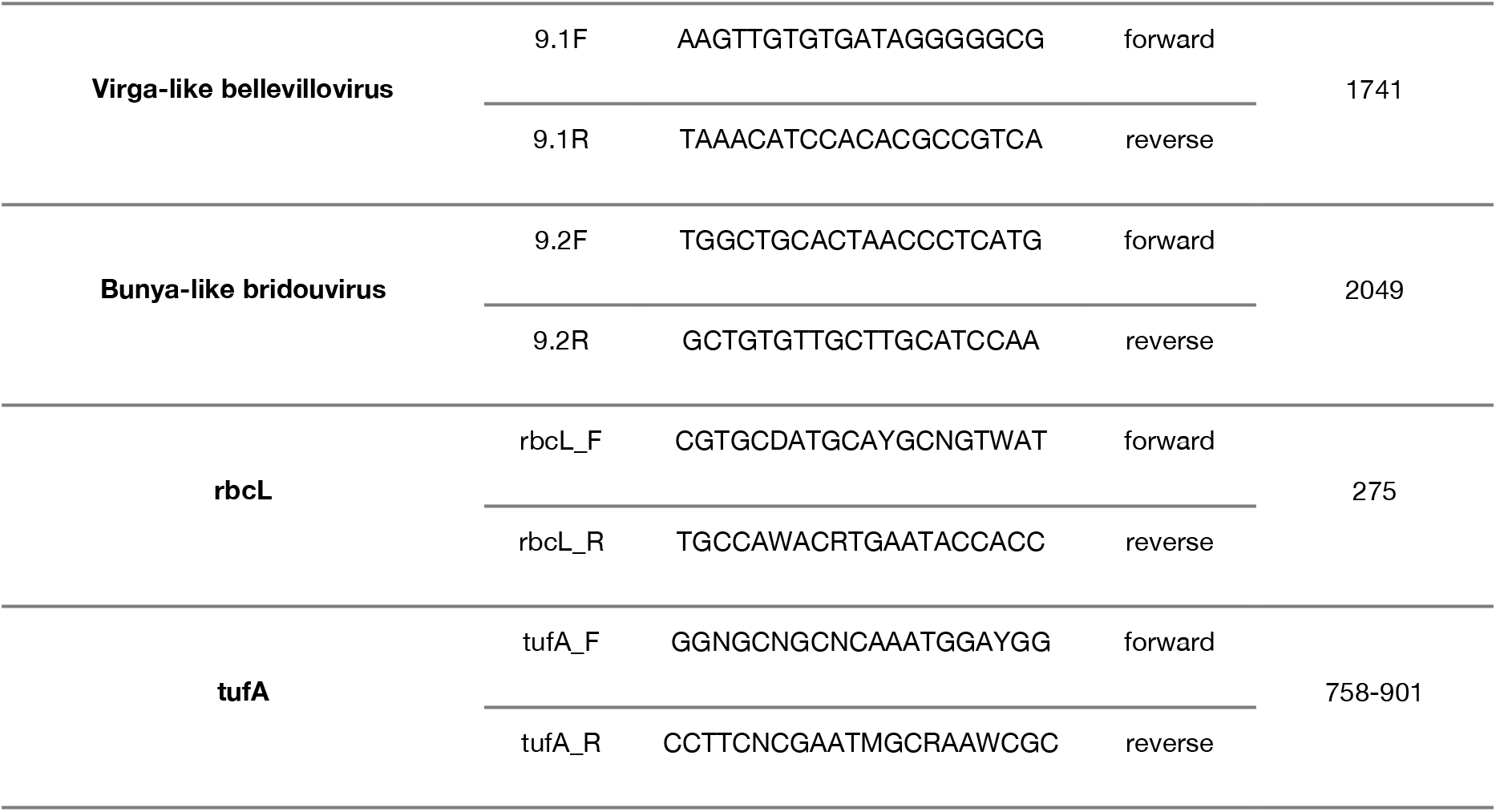
List of primers used in this study. tufA primers have been retrieved from ^76^.

## Notes

### Competing Interest Statement

The authors have declared no competing interest.

## REFERENCES

1. Krupovic M, Prangishvili D, Hendrix RW, Bamford DH. Genomics of bacterial and archaeal Viruses: dynamics within the prokaryotic virosphere. Microbiol Mol Biol Rev 2011; 75: 610–635.

2. Lang AS, Rise ML, Culley AI, Steward GF. RNA viruses in the sea. FEMS Microbiol Rev 2009; 33: 295–323.

3. Krishnamurthy SR, Wang D. Origins and challenges of viral dark matter. Virus Res 2017; 239: 136–142.

4. Zhang Y-Z, Shi M, Holmes EC. Using metagenomics to characterize an expanding virosphere. Cell 2018; 172: 1168–1172.

5. Short SM, Staniewski MA, Chaban Y V., Long AM, Wang D. Diversity of viruses infecting eukaryotic algae. Curr Issues Mol Biol 2020; 39: 29–62.

6. Mihara T, Nishimura Y, Shimizu Y, Nishiyama H, Yoshikawa G, Uehara H et al. Linking virus genomes with host taxonomy. Viruses 2016; 8: 66.

7. Richmond A, Hu Q. Handbook of Microalgal Culture: Applied Phycology and Biotechnology: Second Edition. In: Handbook of Microalgal Culture: Applied Phycology and Biotechnology: Second Edition. 2013, pp 1–719.

8. Mayer JA, Taylor FJR. A virus which lyses the marine nanoflagellate Micromonas pusilla. Nature 1979; 281: 299–301.

9. Brown RM. Algal viruses. Adv Virus Res 1972; 17: 243–277.

10. Illergard K, Ardell DH, Elofsson A. Structure is three to ten times more conserved than sequence—A study of structural response in protein cores. Proteins Struct Funct Bioinforma 2009; 77: 499–508.

11. Bamford DH, Grimes JM, Stuart DI. What does structure tell us about virus evolution ? Curr Opin Struct Biol 2005; 15: 655–663.

12. Chen J, Guo M, Wang X, Liu B. A comprehensive review and comparison of different computational methods for protein remote homology detection. Brief Bioinform 2018; 19: 231–244.

13. Singh J, Saxena RC. An Introduction to Microalgae: Diversity and Significance. In: Handbook of Marine Microalgae. Academic Press, 2015, pp 11–24.

14. Archibald JM. The evolution of algae by secondary and tertiary endosymbiosis. Adv Bot Res 2012; 64: 87–118.

15. Burki F, Roger AJ, Brown MW, Simpson AGB. The new tree of eukaryotes. Trends Ecol Evol 2020; 35: 43–55.

16. Guiry MD. How many species of algae are there? J. Phycol. 2012; 48: 1057–1063.

17. Chapman RL. Algae: the world’s most important “plants”—an introduction. Mitig Adapt Strateg Glob Chang 2013; 18: 5–12.

18. Leliaert F, Smith DR, Moreau H, Herron MD, Verbruggen H, Delwiche CF et al. Phylogeny and molecular evolution of the green algae. CRC Crit Rev Plant Sci 2012; 31: 1–46.

19. Urayama SI, Takaki Y, Nunoura T. FLDS: A comprehensive DSRNA sequencing method for intracellular RNA virus surveillance. Microbes Environ 2016; 31: 33–40.

20. Mushegian A, Shipunov A, Elena SF. Changes in the composition of the RNA virome mark evolutionary transitions in green plants. BMC Biol 2016; 14: 68.

21. Jackson C, Knoll AH, Chan CX, Verbruggen H. Plastid phylogenomics with broad taxon sampling further elucidates the distinct evolutionary origins and timing of secondary green plastids. Sci Rep 2018; 8: 1–10.

22. Fuhrman JA. Marine viruses and their biogeochemical and ecological effects. Nature 1999; 399: 541–548.

23. Forterre P, Prangishvili D. The major role of viruses in cellular evolution: Facts and hypotheses. Curr Opin Virol 2013; 3: 558–565.

24. Short SM, Staniewski MA, Chaban Y V, Long AM, Wang D. Diversity of viruses infecting eukaryotic algae. Curr Issues Mol Biol 2020; 39: 29–62.

25. Cortona A Del, Jackson CJ, Bucchini F, Van Bel M, D’hondt S, Skaloud P et al. Neoproterozoic origin and multiple transitions to macroscopic growth in green seaweeds. Proc Natl Acad Sci USA 2020; 117: 2551–2559.

26. Cocquyt E, Verbruggen H, Leliaert F, Clerck O De. Evolution and cytological diversification of the green seaweeds (Ulvophyceae). Mol Biol Evol 2010; 27: 2052–2061.

27. Wolf YI, Kazlauskas D, Iranzo J, Lucía-Sanz A, Kuhn JH, Krupovic M et al. Origins and evolution of the global RNA virome. mBio 2018; 9: e02329–18.

28. Koga R, Horiuchi H, Fukuhara T. Double-stranded RNA replicons associated with chloroplasts of a green alga, Bryopsis cinicola. Plant Mol Biol 2003; 51: 991–999.

29. Marcelino VR, Clausen PTLC, Buchmann JP, Wille M, Iredell JR, Meyer W et al. CCMetagen: comprehensive and accurate identification of eukaryotes and prokaryotes in metagenomic data. Genome Biol 2020; 21: 103.

30. Sabanadzovic S, Valverde RA, Brown JK, Martin RR, Tzanetakis IE. Southern tomato virus: The link between the families *Totiviridae* and *Partitiviridae*. Virus Res 2009; 140: 130–137.

31. Koga R, Fukuhara T, Nitta T. Molecular characterization of a single mitochondria-associated double-stranded RNA in the green alga Bryopsis. Plant Mol Biol 1998; 36: 717–724.

32. Bao Y, Chetvernin V, Tatusova T. Improvements to pairwise sequence comparison (PASC): a genome-based web tool for virus classification. Arch Virol 2014; 159: 3293–3304.

33. Atkins JF, Loughran G, Bhatt PR, Firth AE, Baranov P V. Ribosomal frameshifting and transcriptional slippage: From genetic steganography and cryptography to adventitious use. Nucleic Acids Res 2016; 44: 7007–7078.

34. Krupovic M, Dolja V V, Koonin E V. Plant viruses of the Amalgaviridae family evolved via recombination between viruses with double-stranded and negative-strand RNA genomes. Biol Direct 2015; 10: 12.

35. Zhang T, Jiang Y, Dong W. A novel monopartite dsRNA virus isolated from the phytopathogenic fungus Ustilaginoidea virens and ancestrally related to a mitochondria-associated dsRNA in the green alga Bryopsis. Virology 2014; 462-463: 227–235.

36. Goh CJ, Park D, Lee JS, Sebastiani F, Hahn Y. identification of a novel plant amalgavirus (*Amalgavirus*, *Amalgaviridae*) genome sequence in *Cistus incanus*. Acta Virol 2018; 62: 122–128.

37. Zhan B, Cao M, Wang K, Wang X, Zhou X. Detection and characterization of cucumis melo cryptic virus, cucumis melo amalgavirus 1, and melon necrotic spot virus in cucumis melo. Viruses 2019; 11: 1–15.

38. Lee JS, Goh CJ, Park D, Hahn Y. Identification of a novel plant RNA virus species of the genus Amalgavirus in the family *Amalgaviridae* from chia (*Salvia hispanica*). Genes and Genomics 2019; 41: 507–514.

39. Roossinck MJ. Lifestyles of plant viruses. Philos Trans R Soc B Biol Sci 2010; 365: 1899–1905.

40. Nibert ML, Ghabrial SA, Maiss E, Lesker T, Vainio EJ, Jiang D et al. Taxonomic reorganization of family *Partitiviridae* and other recent progress in partitivirus research. Virus Res 2014; 188: 128–141.

41. Shi M, Lin X-D, Tian J-H, Chen L-J, Chen X, Li C-X et al. Redefining the invertebrate RNA virosphere. Nature 2016; 540: 539–543.

42. Polischuk V, Budzanivska I, Shevchenko T, Oliynik S. Evidence for plant viruses in the region of Argentina Islands, Antarctica. FEMS Microbiol Ecol 2007; 59: 409–417.

43. Petrzik K, Vondrák J, Kvíderová J, Lukavský J. Platinum anniversary: virus and lichen alga together more than 70 years. PLoS One 2015; 10: e0120768.

44. Nibert ML, Vong M, Fugate KK, Debat HJ. Evidence for contemporary plant mitoviruses. Virology 2018; 518: 14–24.

45. Roossinck MJ. Evolutionary and ecological links between plant and fungal viruses. New Phytol 2019; 221: 86–92.

46. Bonfante P. Algae and fungi move from the past to the future. eLife 2019; 8: e49448.

47. Ricci F, Rossetto Marcelino V, Blackall LL, Kühl M, Medina M, Verbruggen H. Beneath the surface: Community assembly and functions of the coral skeleton microbiome. Microbiome 2019; 7: 1–10.

48. C. R, J. R. Fungi and Their Role in Corals and Coral Reef Ecosystems. In: Raghukumar C. (ed). Biology of Marine Fungi. Springer, Berlin, Heidelberg, 2012, pp 89–113.

49. Marcelino VR, Verbruggen H. Multi-marker metabarcoding of coral skeletons reveals a rich microbiome and diverse evolutionary origins of endolithic algae. Sci Rep 2016; 6: 31508.

50. Nerva L, Vigani G, Di Silvestre D, Ciuffo M, Forgia M, Chitarra W et al. Biological and molecular characterization of Chenopodium quinoa mitovirus 1 reveals a distinct small RNA response compared to those of cytoplasmic RNA viruses. J Virol 2019; 93: e01998–18.

51. Charon J, Grigg MJ, Eden JS, Piera KA, Rana H, William T et al. Novel RNA viruses associated with *Plasmodium vivax* in human malaria and *Leucocytozoon* parasites in avian disease. PLoS Pathog 2019; 15: e1008216.

52. Zangger H, Ronet C, Desponds C, Kuhlmann FM, Robinson J, Hartley M-A et al. Detection of Leishmania RNA virus in Leishmania parasites. PLoS Negl Trop Dis 2013; 7: e2006.

53. Grybchuk D, Akopyants NS, Kostygov AY, Konovalovas A, Lye L-F, Dobson DE et al. Viral discovery and diversity in trypanosomatid protozoa with a focus on relatives of the human parasite *Leishmania*. Proc Natl Acad Sci 2018; 115: E506–E515.

54. Cai G, Myers K, Fry WE, Hillman BI. A member of the virus family *Narnaviridae* from the plant pathogenic oomycete *Phytophthora infestans*. Arch Virol 2012; 157: 165–169.

55. Yoon HS, Hackett JD, Ciniglia C, Pinto G, Bhattacharya D. A molecular timeline for the origin of photosynthetic eukaryotes. Mol Biol Evol 2004; 21: 809–818.

56. Parfrey LW, Lahr DJG, Knoll AH, Katz LA. Estimating the timing of early eukaryotic diversification with multigene molecular clocks. Proc Natl Acad Sci U S A 2011; 108: 13624–13629.

57. Burki F, Keeling PJ. Rhizaria. Curr Biol 2014; 24: R103–R107.

58. Gould SB. Algae’s complex origins. Nature 2012; 492: 46–48.

59. Wetherbee R, Rossetto Marcelino V, Costa JF, Grant B, Crawford S, Waller RF et al. A new marine prasinophyte genus alternates between a flagellate and a dominant benthic stage with microrhizoids for adhesion. J Phycol 2019; 55: 1210–1225.

60. Wetherbee R, Verbruggen H. *Kraftionema allantoideum*, a new genus and family of Ulotrichales (Chlorophyta) adapted for survival in high intertidal pools. J Phycol 2016; 52: 704–715.

61. Bolger AM, Lohse M, Usadel B. Trimmomatic: a flexible trimmer for Illumina sequence data. Bioinformatics 2014; 30: 2114–2120.

62. Quast C, Pruesse E, Yilmaz P, Gerken J, Schweer T, Yarza P et al. The SILVA ribosomal RNA gene database project: improved data processing and web-based tools. Nucleic Acids Res 2013; 41: D590–6.

63. Grabherr MG, Haas BJ, Yassour M, Levin JZ, Thompson DA, Amit I et al. Full-length transcriptome assembly from RNA-Seq data without a reference genome. Nat Biotechnol 2011; 29: 644–652.

64. Li B, Dewey CN. RSEM: accurate transcript quantification from RNA-Seq data with or without a reference genome. BMC Bioinformatics 2011; 12: 323.

65. Buchfink B, Xie C, Huson DH. Fast and sensitive protein alignment using DIAMOND. Nat Methods 2015; 12: 59–60.

66. El-Gebali S, Mistry J, Bateman A, Eddy SR, Luciani A, Potter SC et al. The Pfam protein families database in 2019. Nucleic Acids Res 2019; 47: D427–D432.

67. Eddy SR. Accelerated profile HMM searches. PLoS Comput Biol 2011; 7: e1002195.

68. Kelley LA, Mezulis S, Yates CM, Wass MN, Sternberg MJE. The Phyre2 web portal for protein modeling, prediction and analysis. Nat Protoc 2015; 10: 845–858.

69. Remmert M, Biegert A, Hauser A, Söding J. HHblits: lightning-fast iterative protein sequence searching by HMM-HMM alignment. Nat Methods 2012; 9: 173–175.

70. Soding J. Protein homology detection by HMM-HMM comparison. Bioinformatics 2005; 21: 951–960.

71. Kearse M, Moir R, Wilson A, Stones-Havas S, Cheung M, Sturrock S et al. Geneious Basic: An integrated and extendable desktop software platform for the organization and analysis of sequence data. Bioinformatics 2012; 28: 1647–1649.

72. Clausen PTLC, Aarestrup FM, Lund O. Rapid and precise alignment of raw reads against redundant databases with KMA. BMC Bioinformatics 2018; 19: 307.

73. Katoh K, Standley DM. MAFFT Multiple sequence alignment software version 7: Improvements in performance and usability. Mol Biol Evol 2013; 30: 772–780.

74. Nguyen L-T, Schmidt HA, von Haeseler A, Minh BQ. IQ-TREE: A fast and effective stochastic algorithm for estimating maximum-likelihood phylogenies. Mol Biol Evol 2015; 32: 268–274.

75. Kalyaanamoorthy S, Minh BQ, Wong TKF, von Haeseler A, Jermiin LS. ModelFinder: fast model selection for accurate phylogenetic estimates. Nat Methods 2017; 14: 587–589.

76. Vieira HH, Bagatini IL, Guinart CM, Vieira AAH, Vieira HH, Bagatini IL et al. tufA gene as molecular marker for freshwater Chlorophyceae. Algae 2016; 31: 155–165.

77. Strassert JFH, Jamy M, Mylnikov AP, Tikhonenkov D V, Burki F. New phylogenomic analysis of the enigmatic phylum telonemia further resolves the eukaryote tree of Llfe. Mol Biol Evol 2019; 36: 757–765.

