## Supporting information for "Meta-transcriptomic detection of diverse and divergent RNA viruses in green and chlorarachniophyte algae"

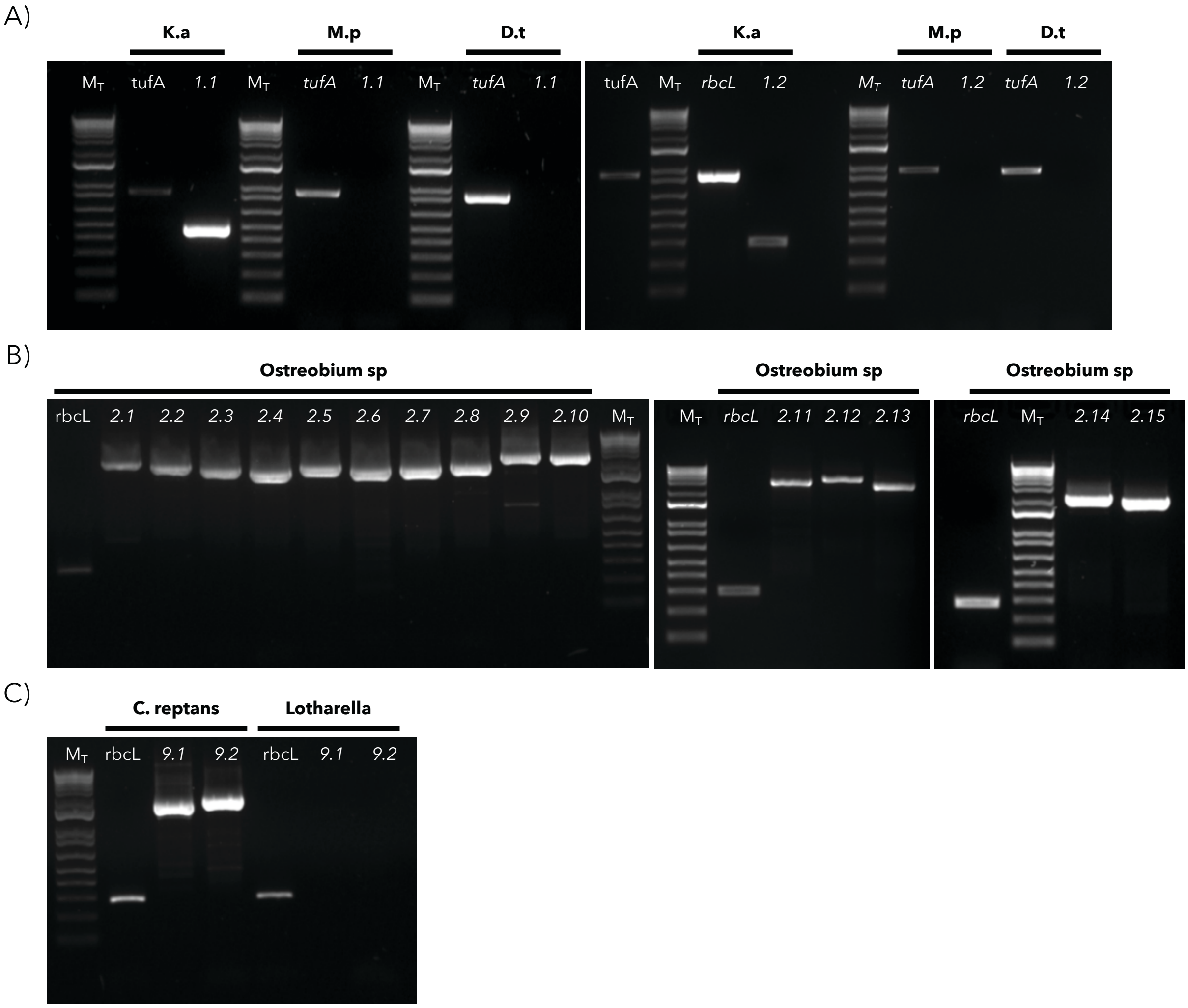


**Fig S1**

**Fig S2** is inserted individually below for purposes of clarity.

**Fig S3** is inserted individually below for purposes of clarity.

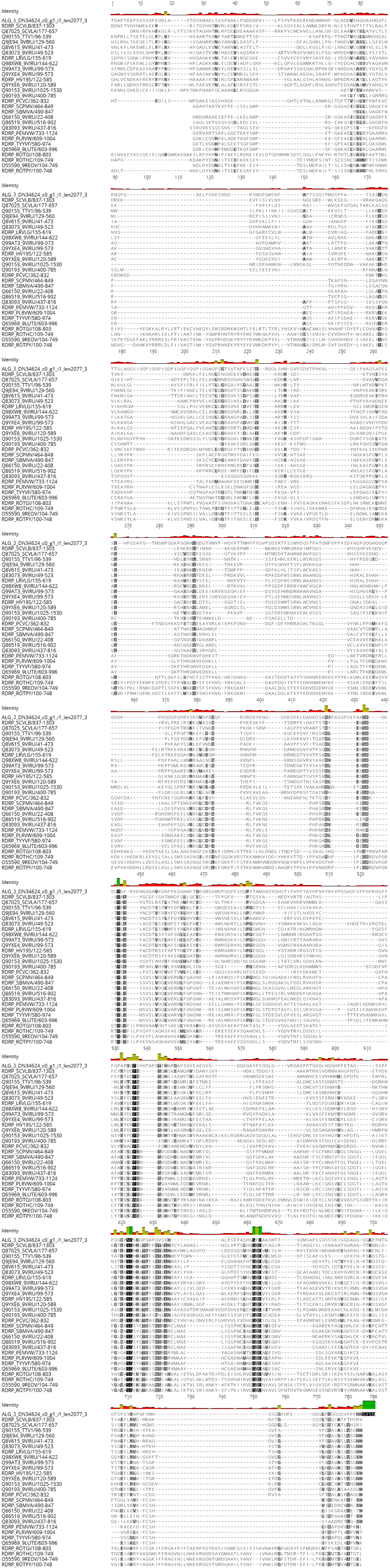

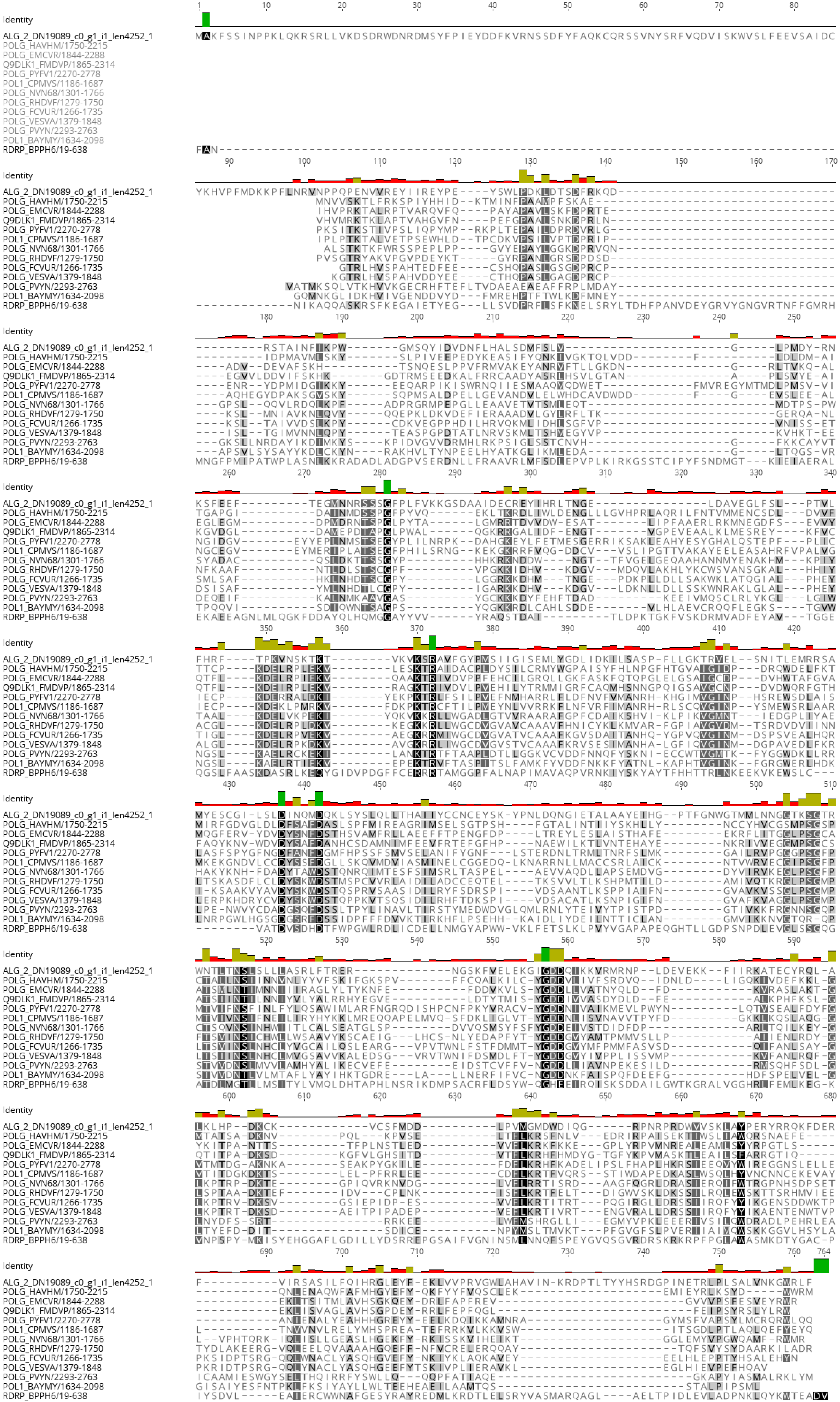
Identity
ALG_2_DN19089_c0_g1_i1_len4252_1
POLG_HAVHM/1750-2215
POLG_EMCVR/1844-2288
Q9DLK1_FMDVP/1865-2314
POLG_PYFV1/2270-2778
POL1_CPMVS/1186-1687
POLG_NVN68/1301-1766
POLG_RHDVF/1279-1750
POLG_FCVUR/1266-1735
POLG_VESVA/1379-1848
POLG_PVYN/2293-2763
POL1_BAYMY/1634-2098
RDRP_BPPH6/19-638

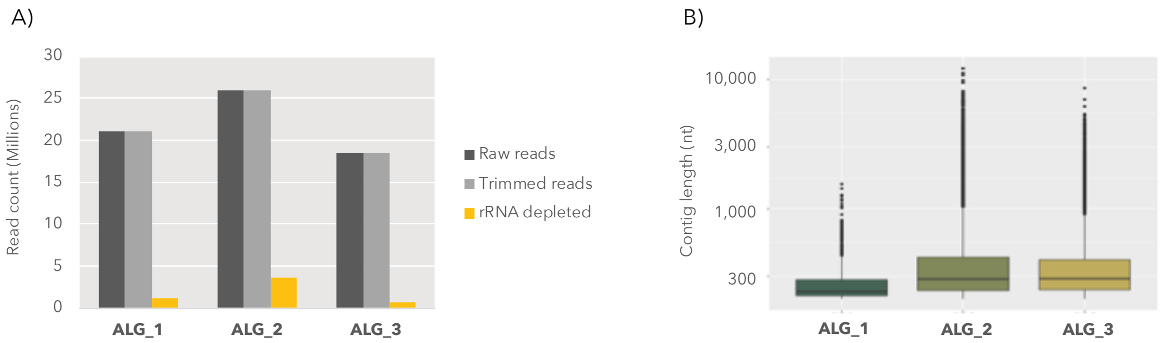
A)
Read count (Millions)
0
5
10
15
20
25
30
ALG_1
ALG_2
ALG_3
Raw reads
Trimmed reads
rRNA depleted
B)
Contig length (nt)
300
1,000
3,000
10,000
ALG_1
ALG_2
ALG_3

**Fig S4**

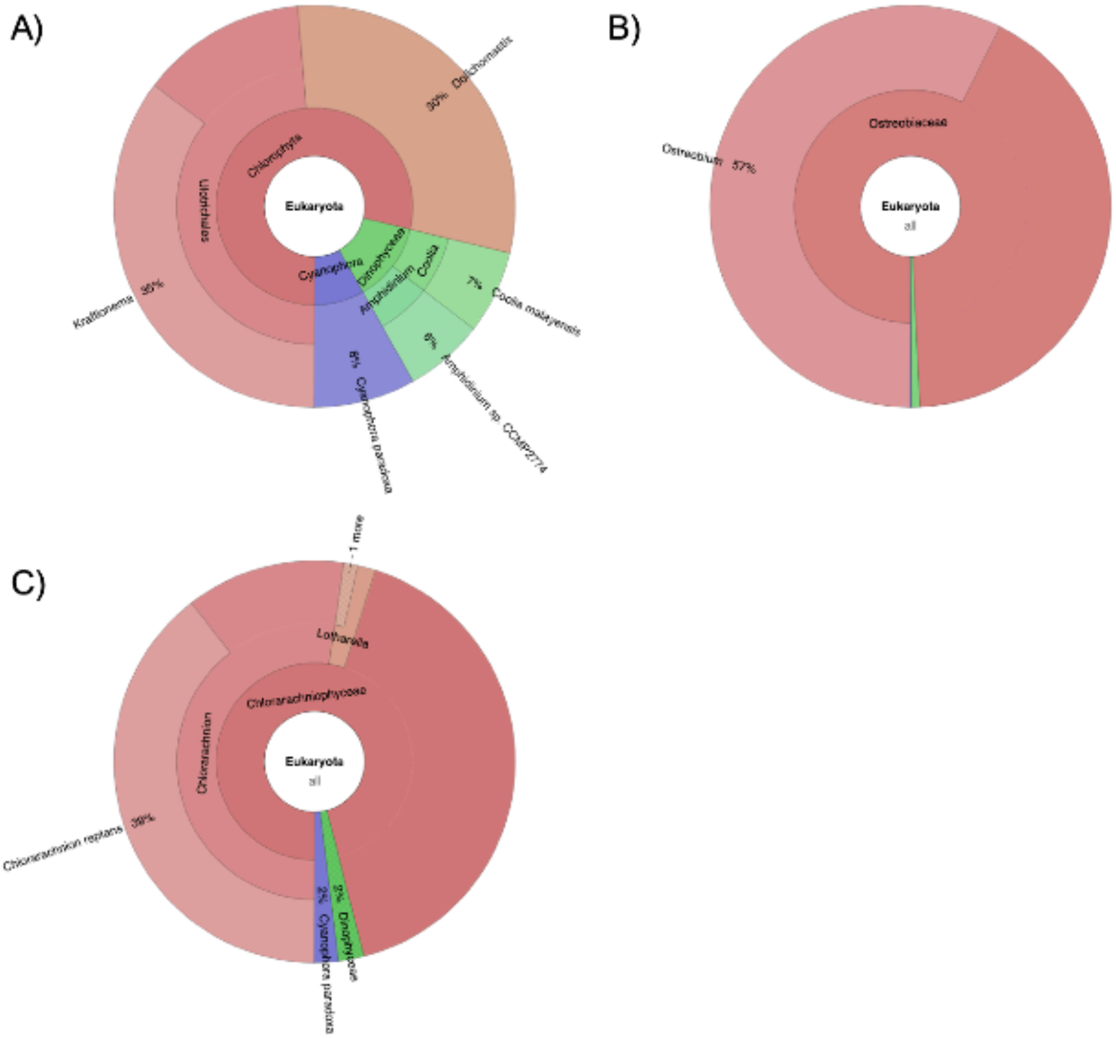
A)
Eukaryota
Chlorophyta
Ulotrichales
30% Dolichomastix
Kraftionema 35%
Cyanophora
Dinophyceae
Amphidinium
Coolia
7% Coolia malayensis
6% Amphidinium sp. CCMP2774
8% Cyanophora paradoxa
B)
Ostreobiaceae
Ostreobium 57%
Eukaryota
all
C)
1 more
Lotharella
Chlorarachniophyceae
Chlorarachnion
Eukaryota
all
Chlorarachnion reptans 39%
2% Cyanophora paradoxa
2% Dinophyceae

**Fig S5**

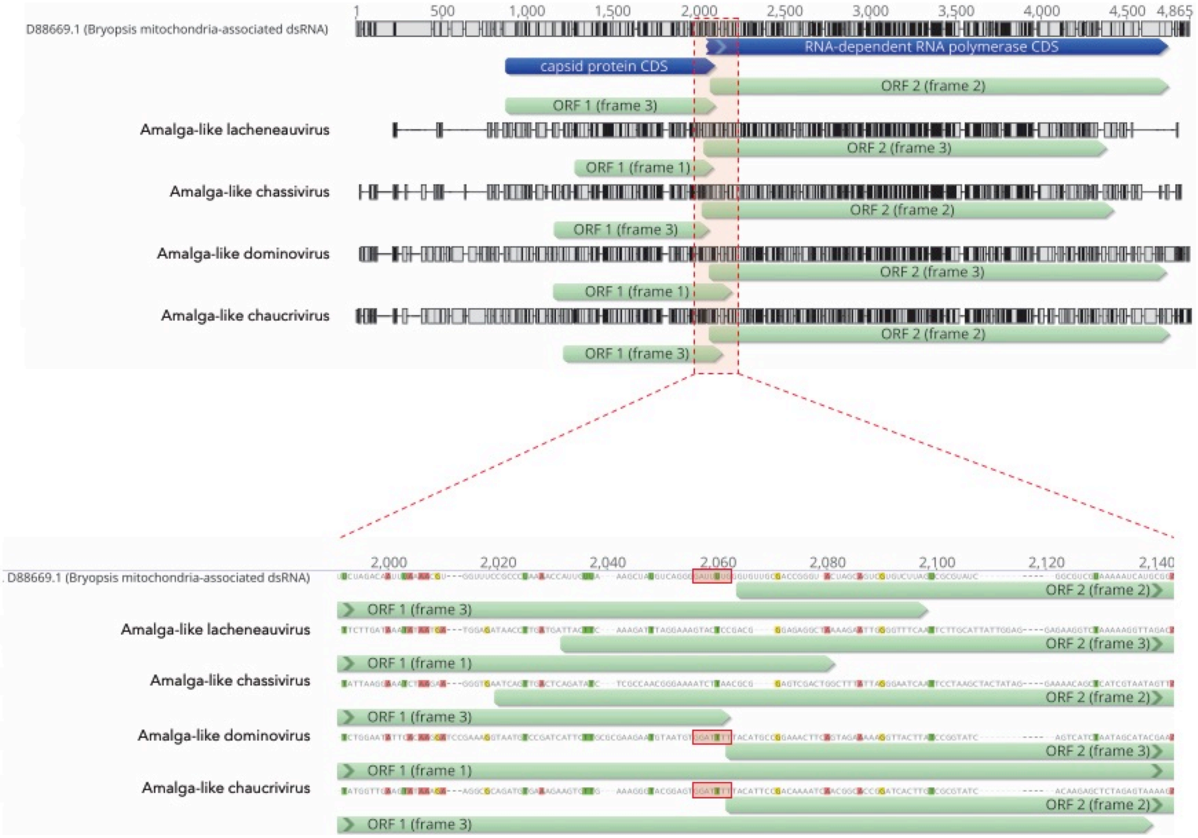

D88669.1 (Bryopsis mitochondria-associated dsRNA)
RNA-dependent RNA polymerase CDS
capsid protein CDS
ORF 2 (frame 2)
ORF 1 (frame 3)
Amalga-like lacheneauvirus
ORF 2 (frame 3)
ORF 1 (frame 1)
Amalga-like chassivirus
ORF 2 (frame 2)
ORF 1 (frame 3)
Amalga-like dominovirus
ORF 2 (frame 3)
ORF 1 (frame 1)
Amalga-like chaucrivirus
ORF 2 (frame 2)
ORF 1 (frame 3)


Fig S6
