## Supplementary figures and images for "Meta-transcriptomic detection of diverse and divergent RNA viruses in green and chlorarachniophyte algae"

### Figure S2

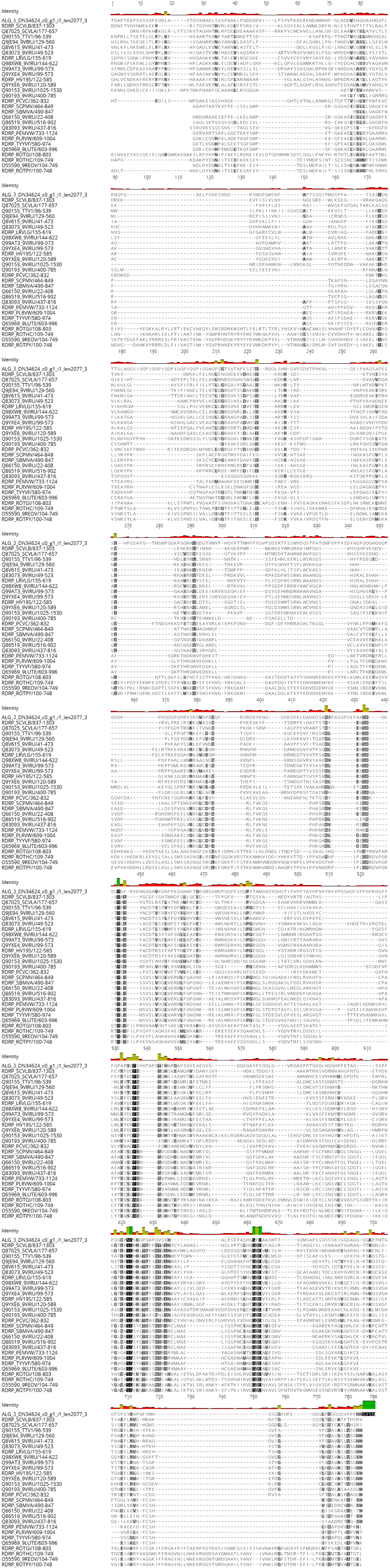

### Figure S3

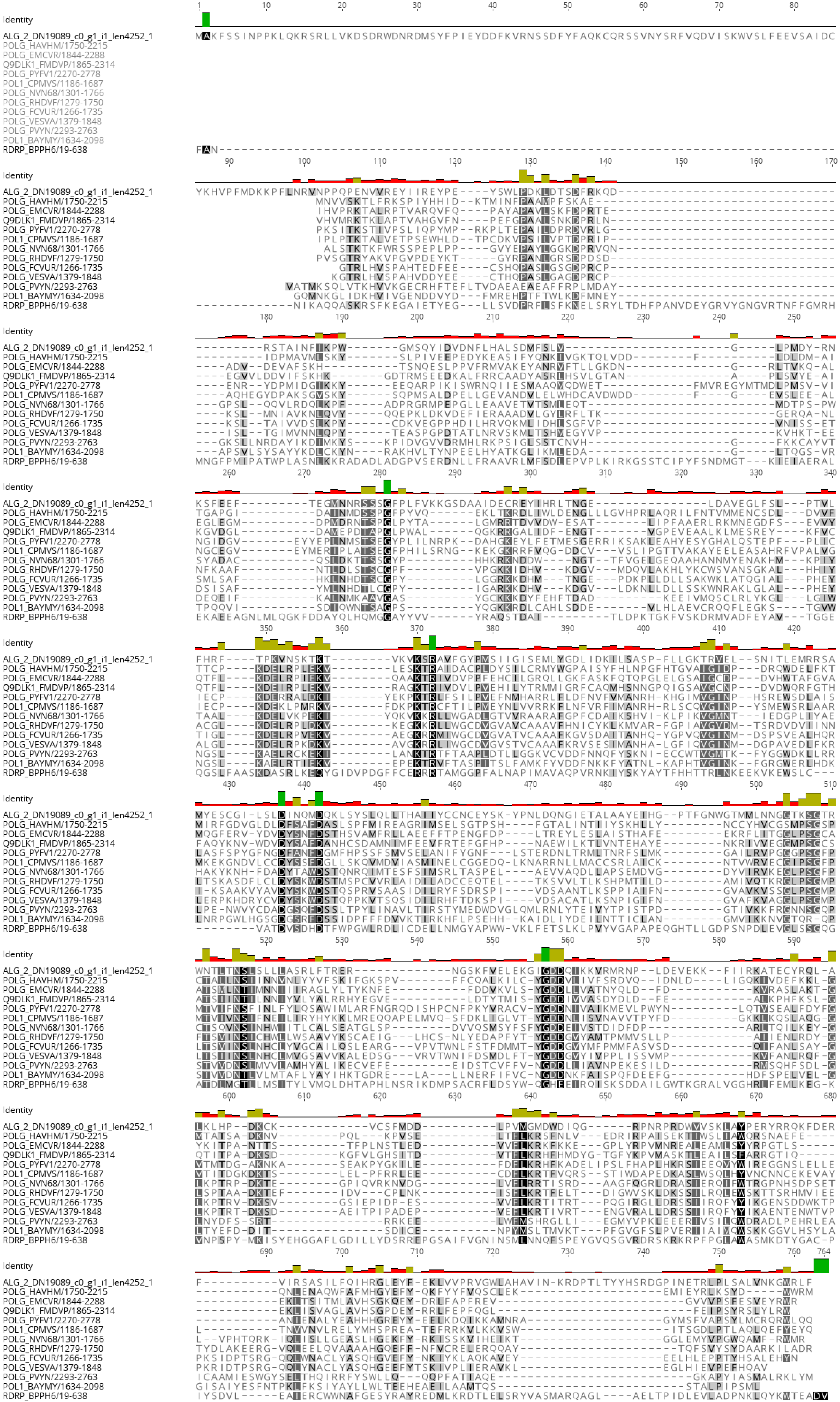
